# SLC35A2 loss of function variants affect glycomic signatures, neuronal fate, and network dynamics

**DOI:** 10.1101/2024.12.27.630524

**Authors:** Dulcie Lai, Paulina Sosicka, Damian J Williams, MaryAnn E Bowyer, Andrew K Ressler, Sarah E Kohrt, Savannah J Muron, Peter B Crino, Hudson H Freeze, Michael J Boland, Erin L Heinzen

## Abstract

*SLC35A2* encodes a UDP-galactose transporter essential for glycosylation of proteins and galactosylation of lipids and glycosaminoglycans. Germline genetic *SLC35A2* variants have been identified in congenital disorders of glycosylation and somatic *SLC35A2* variants have been linked to intractable epilepsy associated with malformations of cortical development. However, the functional consequences of these pathogenic variants on brain development and network integrity remain elusive.

In this study, we use an isogenic human induced pluripotent stem cell-derived neuron model to comprehensively interrogate the functional impact of loss of function variants in *SLC35A2* through the integration of cellular and molecular biology, protein glycosylation analysis, neural network dynamics, and single cell electrophysiology.

We show that loss of function variants in *SLC35A2* result in disrupted glycomic signatures and precocious neurodevelopment, yielding hypoactive, asynchronous neural networks. This aberrant network activity is attributed to an inhibitory/excitatory imbalance as characterization of neural composition revealed preferential differentiation of *SLC35A2* loss of function variants towards the GABAergic fate. Additionally, electrophysiological recordings of synaptic activity reveal a shift in excitatory/inhibitory balance towards increased inhibitory drive, indicating changes occurring specifically at the pre-synaptic terminal.

Our study is the first to provide mechanistic insight regarding the early development and functional connectivity of *SLC35A2* loss of function variant harboring human neurons, providing important groundwork for future exploration of potential therapeutic interventions.

## Introduction

*SLC35A2* (Xp11.23) encodes a Golgi-localized uridine diphosphate (UDP)-galactose transporter likely comprised of 10 transmembrane domains.^1,2^ Functionally, SLC35A2 transports UDP-galactose from the cytosol into the Golgi lumen where it is utilized by glycosyltransferases for the glycosylation of proteins and galactosylation of lipids and glycosaminoglycans.^3^ Cellular glycosylation is fundamentally important for the structural integrity, trafficking, localization, and function of glycoproteins.^3^ In the central nervous system (CNS), cellular glycoproteins mediate critical processes such as axon fate, regeneration, myelination, neural excitability, transmission, and ion channel function^4,5^ Consequently, dysregulated glycosylation may have wide-ranging effects on neurodevelopment, neuron excitability, and network connectivity, which may underlie neurodevelopmental disorders and associated co-morbidities (e.g., epilepsy, intellectual disability, etc.).

*De novo* germline loss-of-function (LOF) genetic variants in *SLC35A2* were first identified in one type of congenital disorder of glycosylation (CDG, OMIM #300896).^2^ CDGs are a group of rare metabolic, multi-system disorders with varying clinical presentations and severity.^6^ Despite the phenotypic heterogeneity, more than 80% of individuals with a CDG present with profound neurological deficits including epilepsy and intellectual disability.^6^ *De novo* germline LOF *SLC35A2* variants were also identified in individuals diagnosed with early onset epileptic encephalopathy (EOEE) that is characterized by intractable seizures and intellectual disability further highlighting an important role of this transporter in neurological function and connectivity.^7–9^

Malformations of cortical development (MCD) account for up to 40% of drug resistant epilepsy.^10^ These structural brain lesions are acquired during the specific cellular phases of neurodevelopment (e.g., proliferation, migration, differentiation, cortical organization), resulting in a spectrum of malformations.^11,12^ *De novo*, post-zygotically acquired (somatic) variants that arise during embryonic brain development have emerged as an important genetic etiology underlying MCDs. Somatic variants in mTOR pathway genes (e.g., *AKT1, AKT3, mTOR, PIK3CA, RHEB, RPS6, DEPDC5, TSC1, TSC2*) have been extensively characterized in focal cortical dysplasia type II (FCDII) and hemimegalencephaly (HMEG), two subtypes of MCDs.^13–22^ However, our lab was the first to identify a non-mTOR-related mechanism, discovering somatic variants in *SLC35A2* associated with focal cortical dysplasia type I (FCDI) and non-lesional focal epilepsy.^23^ This finding has been substantiated by two additional research groups demonstrating that somatic *SLC35A2* variants account for up to 30% of cases presenting with mild MCD (mMCD)/FCDI.^24,25^ Following the initial discovery of somatic *SLC35A2* variants in MCD, the associated neuropathology was later clarified to be a mild malformation of cortical development with oligodendroglial hyperplasia in epilepsy (MOGHE).^26,27^ Despite these significant findings, the role of *SLC35A2* in neurodevelopment and its contribution to the epilepsy phenotype remain unknown.

In this study, we comprehensively interrogate the effect of *SLC35A2* LOF variants on neurodevelopment and neural network connectivity using an induced pluripotent stem cell (iPSC)-derived neuron model. To accomplish this, we used an isogenic model of human iPSCs derived from a healthy male CRISPR/Cas9-edited to harbor a patient identified missense variant (SLC35A2^S304P/Y^) and knockout (SLC35A2^-/Y^) in *SLC35A2*; both predicted and functionally shown to be LOF variants. Directed differentiation using a protocol that generates predominantly dorsal forebrain glutamatergic neurons reveals that *SLC35A2* LOF variants lead to disrupted glycosylation, in both iPSCs and neurons, and dramatically perturb early neurodevelopment. Furthermore, SLC35A2^S304P/Y^ and SLC35A2^-/Y^ neural networks displayed reduced activity, lack of synchronous firing, and increased burst duration compared to the isogenic control. We find that these differences in network phenotypes are attributed to an inhibitory/excitatory imbalance in neural composition that may be in part driven by changes in activity at the presynaptic terminal. This is the first study to investigate the functional effects of *SLC35A2* LOF variants on neurodevelopment and neural network connectivity using an iPSC-derived neuron model, shedding light on the underlying mechanisms. Additionally, we identify a novel role for *SLC35A2* in the determination of cellular fate.

## Materials and methods

### Human induced pluripotent stem cell (iPSC) culture

The control human iPSC line used in this study (713-5) were reprogrammed from fibroblasts obtained from a healthy male, as previously published.^28^ All iPSC lines (control, SLC35A2^S304P/Y^, SLC35A2^-/Y^) were maintained under feeder-free conditions on Geltrex-coated plates (Gibco, A1413302) in TeSR-E8 (Stem Cell Technologies, 05990) at 37°C and 5% CO_2_. Cultures were passaged with 0.5 mM EDTA and were routinely tested for mycoplasma (Bulldog Bio, 25234) and subject to G-band karyotyping analysis (KaryoLogic, Inc., Durham, North Carolina). Refer to Supplementary Materials for CRISPR/Cas9 genome editing methods.

### Immunofluorescent staining

Cells were fixed on 12 mm coverslips with 4% paraformaldehyde for 15 minutes, washed 3x with 1x PBS, and blocked in staining solution (5% normal goat serum, 1% BSA, 0.3% Triton-X) for 15 minutes. Coverslips were then incubated for one hour with primary antibody (diluted in staining solution), washed 3x with 0.2% Triton-X, and incubated for 30 minutes in the dark with secondary antibody diluted 1:1000 in staining solution. Finally, cells were washed 3x with 0.2% Triton-X and 1x with PBS before mounting with Prolong Diamond Antifade Mountant with DAPI (Invitrogen, P36962) and cured overnight in the dark. Primary and secondary antibodies used are listed in Supplementary Table 1. Images were captured using the Nikon Eclipse Ti2 Inverted Fluorescence Microscope and the Nikon NIS-Elements software. Quantification of PAX6+ and ß3-TUBULIN+ neurons were counted manually by a blinded assessor (M.E.B). Neurite length was quantified using NeurphologyJ.^29^

### Western blot

Whole cell lysate was extracted using RIPA Buffer (Thermo Scientific, 89900) containing cOmplete EDTA-free Protease Inhibitor Cocktail (Sigma Aldrich, 11873580001) and PhosSTOP phosphatase inhibitor (Roche, 4906845001). Protein was quantified using the Qubit 4 Fluorometer (Thermo Fisher Scientific). Samples were prepared with 4x Bolt LDS Sample Buffer (Invitrogen, B0007), 10x Bolt Sample Reducing Agent (Invitrogen, B0009), and denatured at 95^c^C for 10 minutes. Protein (10-20 ug) was resolved using Bolt 4-12% Bis-Tris Plus gels (Invitrogen) and transferred to PVDF membranes. Membranes were blocked in 5% Blotting Grade Buffer (Bio-Rad, 1706404) for one hour at room temperature and incubated overnight at 4°C with primary antibody. Membranes were then washed 3x 5 minutes with TBS-T, incubated with secondary antibody for one hour at room temperature and washed again 3×5 minutes with TBS-T. Proteins were visualized using SuperSignal West Pico PLUS Chemiluminescent reagent (Thermo Scientific, 34579) and captured on the iBright FL1000 Imaging System (Thermo Fisher Scientific). To detect SLC35A2 and TRA-1-81, slight modifications to the above method were made. In brief, the membrane was blocked for five hours in 5% milk in PBS-T at room temperature and incubated with primary antibody at 4°C overnight followed by 30 minutes at room temperature. The membrane was washed 6x 5 mins in 1% milk in PBS-T, incubated with secondary antibody for two hours at room temperature, and washed again for 6x 5 min in 1% milk in PBST. Primary and secondary antibodies were diluted using the SignalBoost Immunoreaction Enhancer Kit (Millipore, 407207). Primary and secondary antibody details are found in Supplementary Table 2.

### N-Glycan isolation and analysis

#### Sample preparation

Sample preparation and N-glycan purification are as previously described.^30^ In brief, cells were washed once with DPBS, collected in 500 uL DPBS and centrifuged for 5 min at 300 rcf. The pellet was flash frozen and stored at −80°C. The pellet was then resuspended in 150 uL G7 buffer [50 mM sodium phosphate (pH 7.5), EDTA-free protease inhibitor] containing 40 mM DTT, sonicated twice, incubated for 10 min at 100°C, and sonicated twice more. PNGase F was then added to the lysate and incubated at 37°C overnight.

#### Isolation of N-glycans

The lysate was centrifuged for 20 min at 14,000 rpm. To purify N-glycans, the supernatant was loaded into activated TopTip 10-200uL carbon columns (PolyLC, Inc.).^30^ The column was then washed 2x with 200 uL water and 2x with a solution containing 3% acetonitrile and 0.05% TFA. Glycans were eluted three times with 200 uL (per elution) using a 50% acetonitrile with 0.05% TFA solution and dried using a SpeedVac.

#### Labeling and Hydrophilic Interaction Chromatography (HILIC) purification of N-glycans

N-glycans were labelled with a procainamide hydrochloride (ProA) solution containing the following: 100 uL of a 30%/70% mix of glacial acetic acid/anhydrous DMSO containing 38 mg/mL ProA (Sigma Aldrich, SML2088) and 45 mg/mL 2-picoline borane (Sigma Aldrich, 654213). Dried glycans were dissolved in 5 uL of ProA solution and incubated for 3 hours at 65°C. ProA-labeled glycans were then purified using HILIC TopTip 200 uL solid phase extraction tips (PolyLC Inc., TT200HIL.96). The column was first activated by performing the following: one wash with 200 uL acetonitrile, two washes with 200 uL 0.1% TFA, and three washes with 200 uL of 89% acetonitrile and 1% TFA. The column was then loaded with 200 uL of 89% acetonitrile and 1% TFA and the sample and the flow through collected. Loading and collection of the flow through was repeated two additional times. Finally, the column was washed three times with 200 uL of 89% acetonitrile and 1% TFA and two times with 200 uL of 90% acetonitrile and 0.1% TFA. Samples were eluted three times with 200 uL (per elution) of 0.1% TFA and dried using a SpeedVac.

#### HPLC separation

Labeled N-glycans were separated on an AdvanceBio Glycan Mapping 300Å, 2.1 x 150 mm, 1.8µm UHPLC column (Agilent, 859700-913) with an AdvanceBio Glycan Mapping 300Å, 2.1 x 5 mm, 1.8 µm guard column (Agilent; 821725-905) using 100 mM ammonium formate pH 4.5 (Buffer A) and acetonitrile (Buffer B) on the Vanquish HPLC system (Thermo Fisher Scientific). The column was first equilibrated with acetonitrile for 70 minutes at a flow rate of 0.5 mL/min, followed by an 85%/15% mix of 100 mM ammonium formate (pH 4.5)/acetonitrile for 140 min at 0.25 mL/min, and lastly a 20%/80% mix of 100 mM ammonium formate (pH 4.5)/acetonitrile for 140 min at 0.5 mL/min. One microliter of sample was then injected and separated using the gradient described in Supplementary Table 3. Separation was performed at 40°C. ProA was excited with λ=310 nm and fluorescence was collected at λ=370 nm. Refer to Supplementary Materials for details pertaining to glycan peak assignment.

### Neuronal differentiation

hiPSC were differentiated using a modified version of the dual SMAD inhibition protocol.^28,31^ Cells were dissociated with Accutase and 8.5×10^5^ cells were seeded onto Geltrex-coated 12-well plates in TeSR-E8 supplemented with 10 uM Y-27632 (Hello Bio, HB2297). The following day neural induction was initiated with neural expansion media [Neurobasal Medium (Gibco, 12348017), 1x N-2 (Gibco, 17502048), 1x B-27 (Gibco, 17504044), 1x MEM Non-Essential Amino Acids (NEAA; Gibco, 11140050), 2 mM L-glutamine (Gibco, 25030081)] supplemented with 1 uM Dorsomorphin (Reprocell, 04-0024) and 10 uM SB-431542 (Reprocell, 04-0010-05). Media was changed daily for the following seven days. On day nine through 23, cells were maintained in neural expansion media with daily media changes. An aggregate passage was performed on day 16-18 with 1 mg/mL collagenase dispase (Sigma, 10269638001) and re-plated onto PDL/laminin coated 12-well plates and/or 12 mm glass coverslips. On day 24, neurons were dissociated with papain (Worthington Biochemical, LK003178) and co-cultured with mouse astrocytes on 48-well MEA plates, 12 mm glass coverslips for immunofluorescent staining, 15 mm glass coverslips for electrophysiology, or cryopreserved in neural expansion media supplemented with 10% DMSO. From day 25 onwards, cultures were maintained in neural maturation media [neural expansion media supplemented with 0.2 mM L-ascorbic acid (Tocris, 40-555-0), 20 ng/mL BDNF (R&D Systems, 248-BDB), 20 ng/mL GDNF (R&D Systems, 212-GD), 125 uM dibutyryl-cAMP (Selleck Chemicals, S7858), 10 uM DAPT (Reprocell, 04-0041), 1x Pen/Strep (15140122)] with 50% media changes every other day.

### Multielectrode array (MEA)

Neurons were dissociated at day 24 of differentiation and plated onto PDL-coated 48-well MEA plates (CytoView MEA 48, Axion Biosystems). Each MEA well consists of 16 electrodes arranged in a 4 x 4 grid spaced 350 µM apart. Neurons were plated at a density of 40,000 cells per well along with 10,000 mouse astrocytes (see Supplementary Methods for cortical astrocyte isolation) in neural expansion media supplemented with 10 uM Y-27632, 2% FBS, and 2 ug/mL laminin and plated as a 40 uL droplet. After an hour incubation at room temperature, 500 uL of neural expansion media supplemented with 10 uM Y-27632, 2% FBS, and 2 ug/mL laminin was added per well and maintained in a 5% CO_2_ and 37°C incubator. The following day a 100% media change to neural maturation media was performed followed by 50% media changes 3x per week thereafter. After approximately two weeks, cultures were then maintained in BrainPhys (supplemented with 0.2 mM L-ascorbic acid, 20 ng/mL BDNF, 20 ng/mL GDNF, 125 uM dibutyryl-cAMP, 1x Pen/Strep). MEA recordings were performed on the Axion Maestro Pro (Axion Biosystems) at 37°C and 5% CO_2_ using the AxIS Navigator software. MEA plates were first equilibrated in the apparatus for 5 minutes prior to a 15 minute recording. Activity was recorded 3x per week for two months.

### MEA data analysis

The meaRtools open-source R package was used to extract a total of 70 spike, burst, and network features that are used to characterize neural activity at electrode and network levels.^32^ Data was filtered to remove inactive electrodes and wells. An active electrode detects at least 5 spikes per minute and an active well requires at least 4 active electrodes (25% of all electrodes) per well in more than 50% of the recordings. An adaptive threshold spike detector of 6x the standard deviation (SD) of noise was applied to all recordings.

### Pharmacologic perturbation of neural networks on MEA

Mature neural networks (> 60 days on MEA) were acutely treated with 10 uM of bicuculline (GABA-A antagonist; Selleck Chemicals, S7071), 50 uM CNQX (AMPA-R antagonist; Sigma, C239), or 1 uM TTX (sodium channel blocker; Biotium, 00061). A baseline recording was first taken from the MEA plate. Spent media was transferred out and reserved, leaving 150 uL behind. Drug or vehicle was then spiked into each well and gently mixed to distribute. The treated MEA plate was then equilibrated for 5 minutes followed by a 15 minute recording. Media was then completely aspirated, and wells were rinsed 3x with PBS before being replenished with a mix of 50% spent media with 50% fresh media. MEA plates were incubated for 48 hours to allow full recovery of neural networks prior to treatment with another drug.

### qRT-PCR

Total RNA was extracted using an RNeasy Plus Mini Kit (Qiagen, 74134) and quantified using the Qubit 4 Fluorometer. cDNA was generated from 200-400 ng RNA using the High-Capacity cDNA Reverse Transcription Kit (Applied Biosystems, 4368814). Reactions were prepared using pre-validated TaqMan Gene Expression Assays (ThermoFisher, 4331182) and TaqMan Fast Advanced Master Mix (Applied Biosystems, 4444556) and analyzed using the QuantStudio 6 Flex system (Applied Biosystems). Each sample was run in triplicates per probe and normalized to GAPDH to obtain a ΔCt value. Data is presented as −ΔCt values. Taqman probes are found in Supplementary Table 4.

### Electrophysiology

Electrophysiological recordings were carried out using conventional whole-cell current or voltage clamp methods. 3 x 10^5^ neurons were plated onto 15 mm diameter glass coverslips and cultured for 75 – 90 days. Both recording techniques were performed using a Multiclamp 700B amplifier and a Digidata 1550 digital-to-analogue converter (both from Molecular Devices) at a 10 kHz sample frequency and filtered with a 3 kHz low-pass Bessell filter. Patch pipettes were fabricated with a P-97 pipette puller (Sutter Instruments) using 1.5 mm outer diameter, 1.28 mm inner diameter filamented capillary glass (World Precision Instruments). The external recording solution contained (in mM): NaCl 145, KCl 5, HEPES 10, Glucose 10, CaCl_2_ 2, MgCl_2_ 2. The solution had an osmolality of 300 mOsm. All recordings were carried out at room temperature (21–23 °C).

For assessment of synaptic activity, 100 nM TTX was added to the external recording solution. The pipette solution contained (in mM): cesium methanesulfonate 130, sodium methanesulfonate 10, EGTA 10, CaCl_2_ 1, HEPES 10, TEA-Cl 10, MgATP 5, Na_2_GTP 0.5, QX-314 5, pH 7.2 with CsOH, adjusted to 290 mOsm with sucrose. Whole cell voltage clamp technique was used to record miniature excitatory postsynaptic currents (mEPSCs) and miniature inhibitory postsynaptic currents (mIPSCs) from the same cell by holding the cells at −60 and 0 mV, respectively. Recordings were acquired for a period of 5 minutes. Miniature events were detected offline with Clampfit 10.7 (Molecular Devices) using the template matching function and a minimum threshold of 5 pA. Each event was manually inspected to determine inclusion or rejection in analysis.

Quantification was carried out using custom scripts written for Igor Pro (Wavemetrics, USA) and R v.3 (www.R-project.org).

### Statistics

Statistical analyses were conducted using GraphPad Prism 10. Datasets with three or more groups were compared using one-way ANOVA followed by Tukey’s multiple comparisons. Datasets with three or more groups and two or more independent variables were compared using two-way ANOVA followed by Tukey’s multiple comparisons test. Non-normally distributed data, specifically Figure 7C and 7D, were assessed using the Kruskal-Wallis test, followed by Dunn’s test to correct for multiple comparisons. For MEA, reported p-values are representative of Mann-Whitney U tests that were permuted 1000 times.^32^

### Data availability

The data that support the findings of this study are available from the corresponding author, upon reasonable request.

## Results

### CRISPR/Cas9-edited *SLC35A2* variants lead to aberrant glycosylation

To investigate the effects of variants in X-linked *SLC35A2* on neurodevelopment and network connectivity, we developed an isogenic human iPSC model where control iPSCs, derived from a male without any neurodevelopmental disorders,^28^ were CRISPR/Cas9-edited to harbor an *SLC35A2* missense variant (c.910T>C, p.Ser304Pro; SLC35A2^S304P/Y^; refer to Supplementary Methods for CRISPR/Cas9 editing methods). This patient-identified somatic variant was previously found in a person with intractable non-lesional focal epilepsy with a variant allele fraction (VAF) between 3.3-6.6%.^23^ Since CRISPR/Cas9-editing is prone to introducing indels, we also selected a clone harboring a two base-pair deletion to serve as an *SLC35A2* gene knockout (c.911_912del; SLC35A2^-/Y^). The genotype of CRISPR/Cas9-edited iPSCs were confirmed with Sanger sequencing, and all iPSC lines maintained pluripotent characteristics and normal karyotype (Supplementary Fig. 1 and 2). Interestingly, the TRA-1-81 pluripotent marker was undetected in SLC35A2^S304P/Y^ and SLC35A2^-/Y^ iPSCs using immunofluorescent staining and Western blot (Supplementary Fig. 1, D and E). The TRA-1-81 antibody recognizes a glycan epitope on the protein podocalyxin. Podocalyxin levels were detected with a downward shift in molecular weight in SLC35A2^S304P/Y^ and SLC35A2^-/Y^ iPSCs (Supplementary Fig. 1E) relative to control, consistent with a loss of the glycan modifications. This suggests that the absence of TRA-1-81 staining is likely due to the absence of a glycan epitope, due to loss of SLC35A2 in these iPSC lines. Despite a slight increase in mRNA expression in the SLC35A2^-/Y^ iPSC line, suggesting escape from non-sense mediated decay, neither variant shows significant differences in total *SLC35A2* mRNA levels relative to the isogenic control (Fig. 1A). However, SLC35A2^S304P/Y^ and SLC35A2^-/Y^ variants result in complete loss of protein expression and Golgi localization compared to the control, suggesting altered post-transcriptional mechanisms, or protein instability (Fig. 1, B and C). To confirm the functional impact of loss of SLC35A2 expression on cellular glycoproteins, we performed structural profiling of N-glycans isolated from iPSCs (Fig. 1D). HPLC spectra reveal remarkable differences in N-glycan complexity. Specifically, bi-, tri- and tetra-antennary galactosylated N-glycans are depleted in both SLC35A2^S304P/Y^ and SLC35A2^-/Y^ iPSCs compared to the control. The only galactosylated structures present in SLC35A2^S304P/Y^ and SLC35A2^-/Y^ iPSCs are bi-antennary N-glycans containing a single galactose residue. These findings substantiate previous observations in SLC35A2-deficient cell lines and provide strong evidence that loss of SLC35A2 expression significantly impairs galactosylated N-glycans.^33^

**Figure 1.**
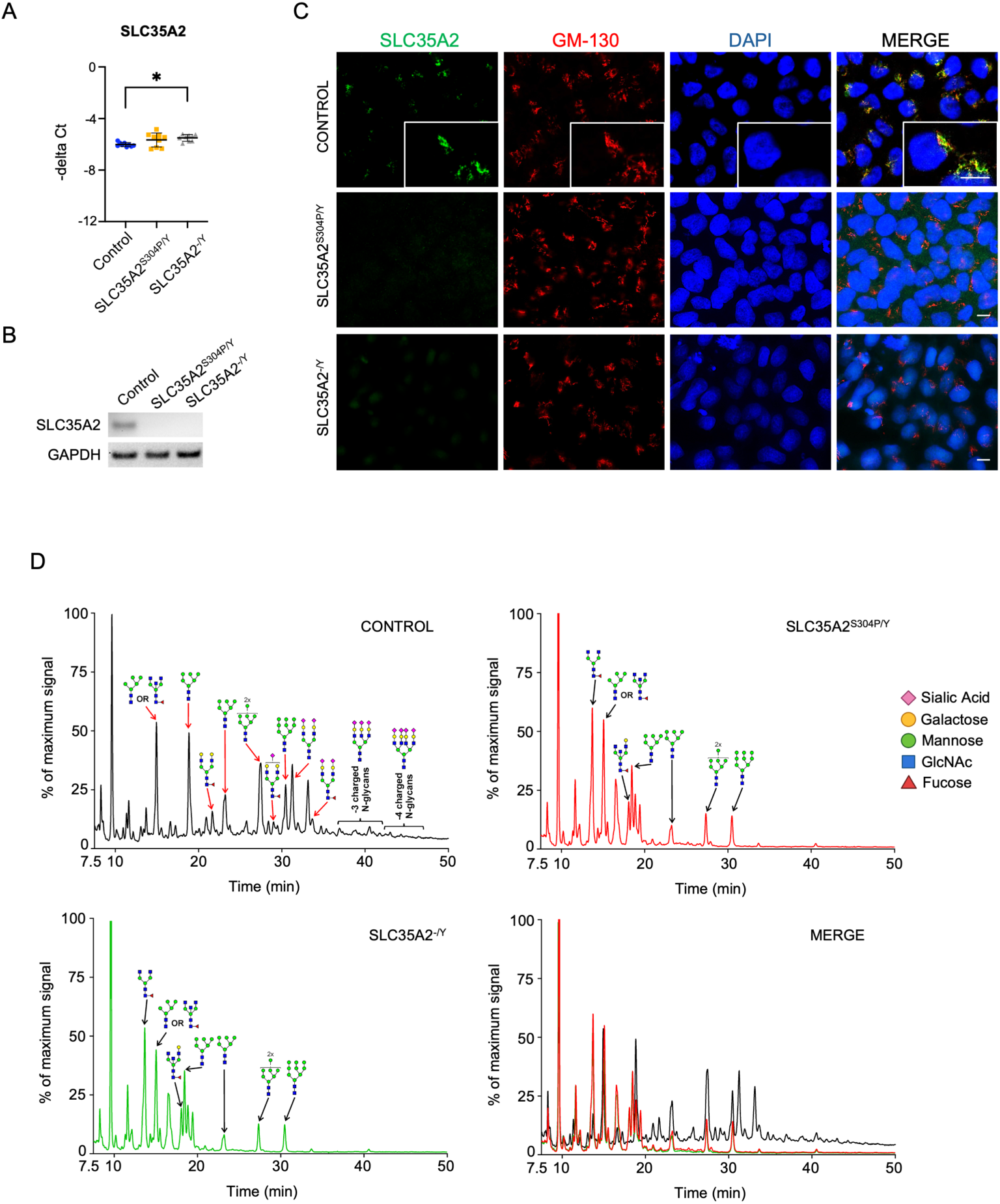
SLC35A2^S304P/Y^ and SLC35A2^-/Y^ encode loss-of-function proteins that result in altered glycosylation in undifferentiated iPSCs. (A) Quantification of SLC35A2 mRNA expression levels by qRT-PCR. Data is representative of three pooled independent replicates each run in triplicate. Each dot represents a single data point. Control is represented as blue circles, SLC35A2^S304P/Y^ as orange squares, and SLC35A2^-/Y^ as grey triangles. Statistics: One-way ANOVA with Tukey’s multiple comparisons. (B) Western blot of SLC35A2 protein expression levels. GAPDH was used as a loading control. Un-cropped, full-length blots are provided in the Supplementary Materials. (C) Localization of SLC35A2 protein (green) with the Golgi apparatus (GM-130; red). Scale bar represents 10 µm. (D) Structural profiling of N-glycans in iPSCs. N-glycans were enzymatically released with PNGase F treatment, labeled with ProA, purified on a HILIC resin and separated using HPLC. HPLC spectra for control, SLC35A2^S304P/Y^ and SLC35A2^-/Y^ were merged in the bottom right panel. The data represents one of two independent replicates.

### Loss of SLC35A2 expression results in early neurogenic phenotypes and sustained altered glycosylation

Glycosylation broadly affects many proteins important for neurite outgrowth, synaptic transmission, ion channel function, etc.^4,5^ Therefore, we hypothesized that LOF variants in *SLC35A2* may affect neurodevelopment and network formation. To study the effects of the variants on these processes we used a method of directed differentiation of iPSCs to dorsal forebrain neurons (Fig. 2A)^31,34^ that closely mimics the temporal nature of neocortical development *in vivo*. Neural induction efficiency was >80% for all three lines as quantified by PAX6+ cells and the expression of neural stem cell (NESTIN, SOX2) and neuroepithelial markers (ZO-1) (Supplementary Fig. 3, A, B and E). Importantly, we also detect altered glycosylation in day 30 cultures, which contain a mixture of NPCs and immature neurons, revealing a reduction of complex N-glycan structures in SLC35A2^S304P/Y^ and SLC35A2^-/Y^ neurons compared to the control (Fig. 2B).

**Figure 2.**
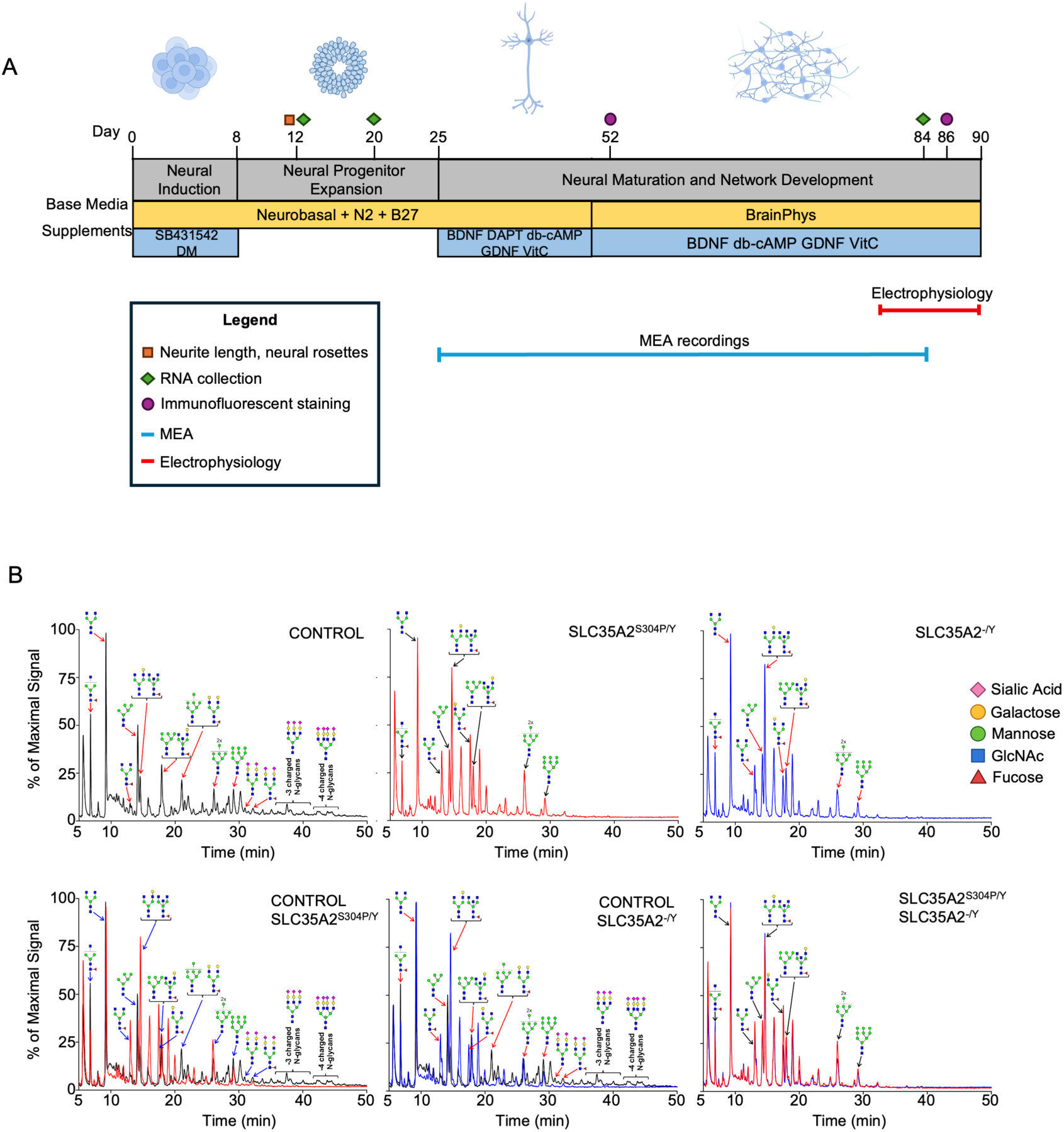
Altered N-glycosylation profiles during neuronal differentiation. **(A)** Experimental workflow depicting directed differentiation protocol and experimental timepoints. Colored icons indicate timepoints at which experiments were conducted. Early neurodevelopmental phenotypes (neurite length and neural rosettes) were assessed at day 12 (orange square). RNA was collected at days 12, 20 and 84 (green diamond). Immunofluorescent staining of neurons was performed at day 52 and 86 (purple circle). MEA recordings were taken following plating at ∼day 24 for two months as indicated by the blue bar. Electrophysiology recordings were performed between day 75-90, as represented by the red bar. **(B)** Structural profiling of N-glycans in cultures at day 30 of differentiation. N-glycans were enzymatically released with PNGase F treatment, labeled with ProA, purified on a HILIC resin and separated using HPLC. HPLC spectra on the top row represents traces for each genotype as indicated. HPLC spectra on the bottom row represent merged traces with the specific genotypes as indicated. The data shown represents one of three independent replicates.

To investigate the effect of *SLC35A2* LOF variants on neurodevelopment, cultures at day 4, 8, and 12 of differentiation were probed for neural rosettes (PAX6, ZO-1) and immature neurons (ß3-TUBULIN). Relative to the isogenic control, defined ZO-1+ apical lumens, a distinguishing characteristic of neural rosettes, and immature neurons were detected much earlier in *SLC35A2* LOF cultures, indicating accelerated neurodevelopment (Fig. 3, A and B; Supplementary Fig. 4 and Fig. 5). For example, day 12 cultures from the isogenic control still retained ZO-1+ lattice-like staining characteristic of adherens and tight junctions among neuroepithelial cells, however the SLC35A2 cultures possessed well defined ZO-1+ lumens. It is worth noting that the classic radial organization of neural progenitors around the central lumen in rosettes appears to be disrupted in the SLC35A2 PAX6+ cells. Immature neurons (ß3-TUBULIN+) were also more developed in SLC35A2^S304P/Y^ and SLC35A2^-/Y^ at day 12 as demonstrated by longer neurite projections compared to the control (Fig. 3, B and C; Supplementary Fig. 5). This difference in neurite length was no longer apparent by day 20 suggesting that the variants have a greater impact on early neurodevelopment (Supplementary Fig. 3C). Interestingly, despite no difference in neurite length at day 20, we did observe a >1.5-fold increase in the number of neurons in SLC35A2^S304P/Y^ and SLC35A2^-/Y^ cultures compared to the control (Supplementary Fig. 3D). This suggests earlier terminal differentiation, which is supported by the precocious rosette formation and enhanced early neurogenesis observed in the *SLC35A2* mutants.

**Figure 3.**
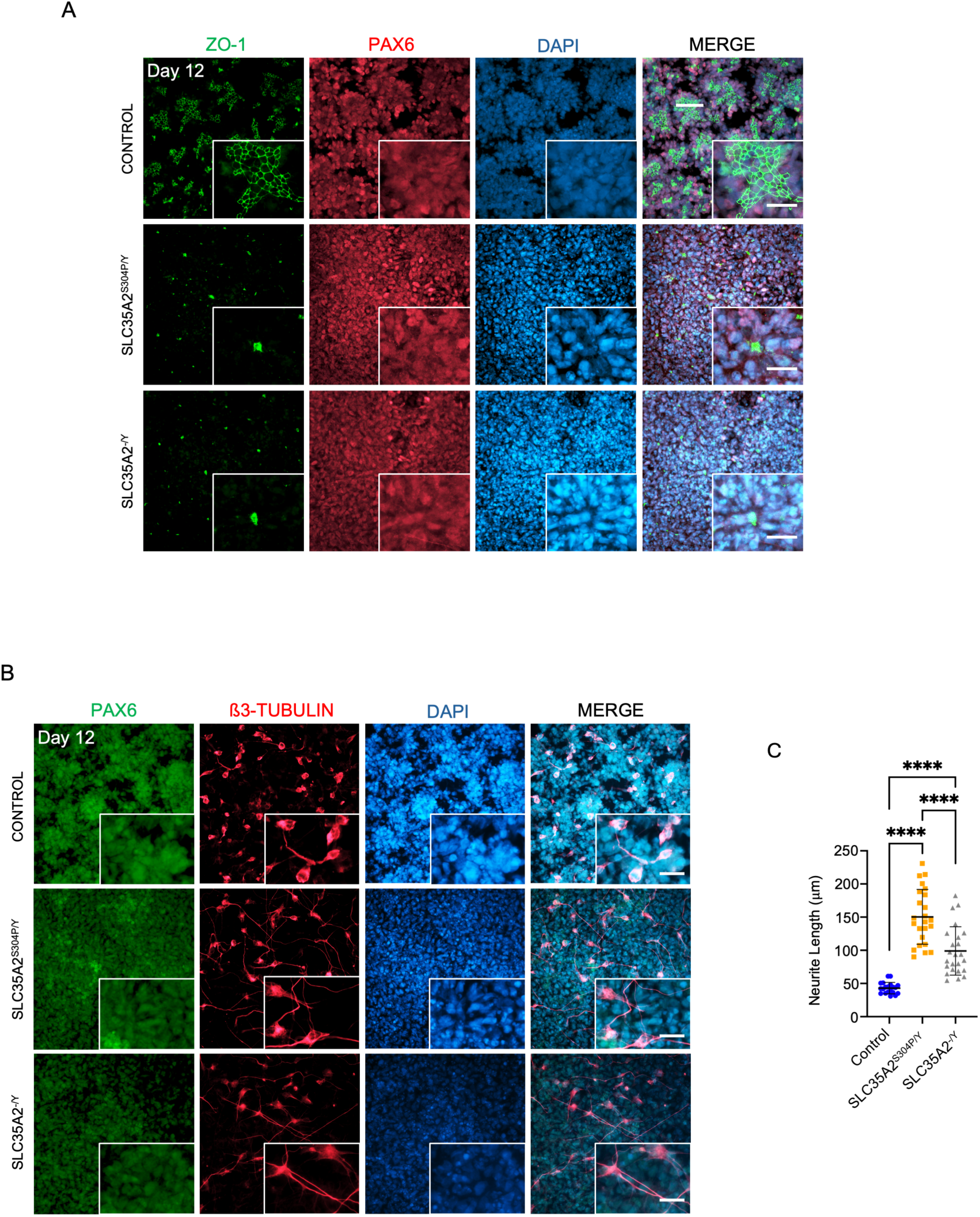
SLC35A2^S304P/Y^ and SLC35A2^-/Y^ promote early neurogenesis. **(A)** Immunofluorescent staining for neural rosettes using the apical lumen marker, ZO-1 (green), and neural progenitor marker, PAX6 (red), at day 12 of differentiation. Nuclei were counterstained with DAPI (blue). Scalebar within the inset represents 20 µm. **(B)** Immunofluorescent staining of day 12 neural cultures for PAX6 (green) and ß3-TUBULIN (red). Nuclei were counterstained with DAPI (blue). Scalebar within the inset represents 20 µm. **(C)** Quantification of neurite length. One-way ANOVA with Tukey’s multiple comparisons. Data is representative of two pooled independent differentiations, with a total of three replicates. Eight regions of interest (ROI) were captured per coverslip. Each dot represents quantification from one ROI. * = P≤0.05; ** = P≤0.01; *** = P≤0.001 **** = P≤0.0001.

**Figure 4.**
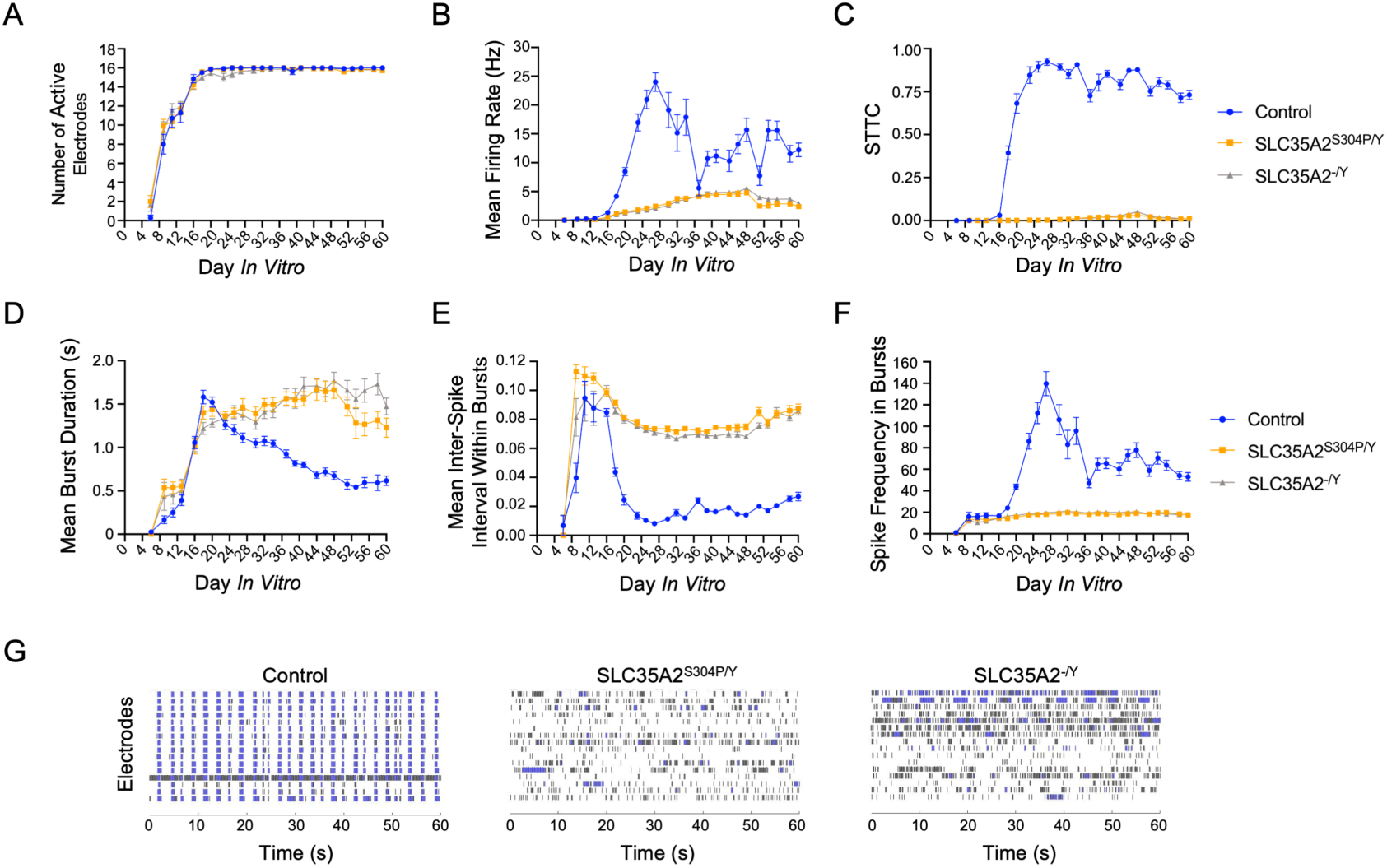
SLC35A2^S304P/Y^ and SLC35A2^-/Y^ generate hypoactive, asynchronous neural networks with prolonged bursting. **A-F)** Graphs depicting MEA features extracted from meaRtools. Each graph contains all recordings for the duration of the experiment. Day *in vitro* (DIV) represents the number of days the neural networks were co-cultured with mouse astrocytes on MEA. Control networks in blue circles, SLC35A2^S304P/Y^ networks in orange squares, and SLC35A2^-/Y^ networks in grey triangles. Data shown is representative of one of four independent differentiations (Refer to Experiment 1 in Supplementary Figures 7 and 8). This differentiation included a technical replicate (A and B), each with 7 wells plated on MEA. The data represents the pooled average of the 14 wells per genotype with error bars representing the standard error. The number of technical replicates and wells plated per Experiment is summarized in Supplementary Table 5. P-value was derived using Mann Whitney test permuted 1000 times (see Methods). **(A)** Number of active electrodes. **(B)** Mean firing rate (MFR) by active electrode. **(C)** Spike train tiling coefficient (STTC). **(D)** Mean burst duration. **(E)** Mean inter-spike interval within bursts. **(F)** Spike frequency within bursts. **(G)** Raster plots from DIV60 (day 84 of differentiation). Each row of the y-axis represents the activity of one of the 16 electrodes. Spikes are represented in black and bursts represented in blue. Plots depict a 60 sec snapshot from a 15 minute recording.

**Figure 5.**
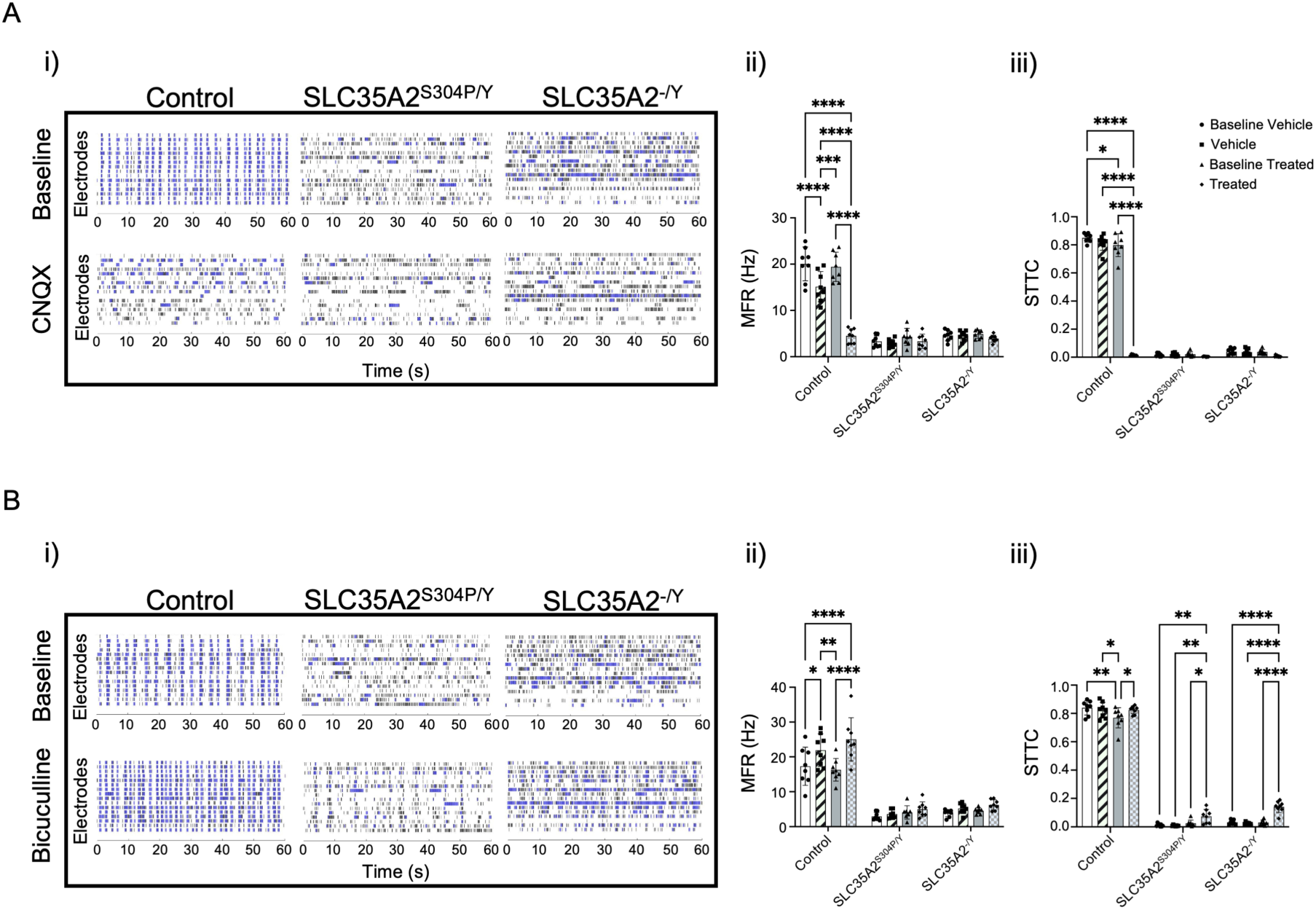
Pharmacologic perturbation of neural networks supports differences in neural composition. A baseline recording of neural networks were first obtained. Respective wells were then treated with vehicle or the appropriate pharmacologic agent. The treated MEA was then allowed to equilibrate for 5 minutes prior to recording. Data is representative of the mean and standard deviation of MEA wells pooled from two independent differentiations. Each dot represents data obtained from a single MEA well. **(A)** Treatment of neural networks with the AMPA/kainate receptor antagonist, CNQX. **(i)** Raster plots depicting a 60 second snapshot of network activity at baseline (top row) and following acute CNQX treatment (bottom row). **(ii)** The effect of CNQX on MFR by active electrode. **(iii)** The effect of CNQX on STTC. **(B)** Treatment of neural networks with the competitive GABA-A receptor antagonist, bicuculline. **(i)** Raster plot depicting a 60 second snapshot of network activity at baseline (top row) and following acute bicuculline treatment (bottom row). **(ii)** The effect of bicuculline on MFR by active electrode. **(iii)** The effect of bicuculline on STTC. Statistics: Two-way ANOVA test with Tukey’s to correct for multiple comparisons. * = P≤0.05; ** = P≤0.01; *** = P≤0.001 **** = P≤0.0001.

### Reduced activity and asynchronous firing in *SLC35A2* LOF neural networks

Since aberrant neural connectivity underlies the epileptic network and clinical epilepsy phenotypes, understanding how *SLC35A2* LOF variants affect neural connectivity and network development is essential to our understanding of how these variants contribute to epileptogenesis. To characterize neural network development and connectivity, neurons at day 24 of differentiation were plated on multiwell multielectrode arrays (MEA). Neurons were co-cultured with primary mouse astrocytes, which are important modulators of neural maturation, synaptogenesis, and long-term culture.^35^ Cultures remained healthy without any observed neuronal aggregation for the duration of the experiment (Supplementary Fig. 6). The number of wells plated per replicate and the number of MEA replicates are summarized in Supplementary Table 5. Network activity was recorded three times per week for two months to capture development and maturation of neural networks. MEA data was longitudinally analyzed using meaRtools, an R package that extracts electrode level [e.g., spikes, bursts, spike train tiling coefficient (STTC), etc.], network level (e.g., network spikes and bursts, etc.) and other (e.g., inter-burst interval, burst duration, etc.) features providing comprehensive analyses of neuronal networks.^32^

MEA recordings captured significant differences in activity and network connectivity between the control, SLC35A2^S304P/Y^, and SLC35A2^-/Y^ harboring neural networks. Overall network activity, represented by mean firing rate (MFR), was consistently higher in the control networks compared to SLC35A2^S304P/Y^ and SLC35A2^-/Y^ despite no differences in the number of active electrodes, suggesting that the loss of SLC35A2 alters either intrinsic excitability or the inhibitory/excitatory imbalance (Fig. 4, A and B). As expected, control networks developed high levels of synchronous firing, represented by the spike train tiling coefficient (STTC) – a pairwise correlation measurement between electrodes – however synchronous activity was virtually absent in SLC35A2^S304P/Y^ and SLC35A2^-/Y^ networks (Fig. 4, C and G).^32,36^

Extracted burst features reveal additional differences in firing behavior. Compared to the control, mean burst duration is prolonged in SLC35A2^S304P/Y^ and SLC35A2^-/Y^ neural networks, corresponding with an increased inter-spike interval (Fig. 4, D and E). Despite the prolonged burst duration, the spike frequency within bursts in SLC35A2^S304P/Y^ and SLC35A2^-/Y^ networks remain significantly lower than the control (Fig. 4F), which is consistent with the overall decrease in MFR. A summary of spike and burst features are found in Supplementary Fig. 7 and 8, raster plots in Supplementary Fig. 9, along with a summary of statistics in Supplementary Tables 6-8 for each MEA replicate.

### Differential response of *SLC35A2* LOF neural networks to CNQX and bicuculline support differences in neural composition

Based on our MEA data, control neural networks possess very distinct dynamics compared to SLC35A2^S304P/Y^ and SLC35A2^-/Y^ networks. To broadly investigate neuronal composition and possible mechanisms underlying network activity, neural networks derived from control and *SLC35A2* LOF variants were acutely probed with bicuculline (GABA_A_ antagonist), cyanquixaline (CNQX; AMPA/kainate receptor antagonist), or tetrodotoxin (TTX; sodium channel blocker).

Control networks treated with CNQX reduced MFR and completely abolished synchronous firing with no effect on SLC35A2^S304P/Y^ and SLC35A2^-/Y^ networks (Fig. 5A). Conversely, bicuculline increased both MFR and synchronous activity in control networks (Fig. 5B). Although bicuculline had no effect on MFR in SLC35A2^S304P/Y^ and SLC35A2^-/Y^, it partially, yet significantly, restored synchronous firing in SLC35A2^S304P/Y^ and SLC35A2^-/Y^ networks (Fig. 5B).

Given the differential responses in firing and network dynamics observed between control and *SLC35A2* LOF variant harboring networks when treated with pharmacologic agents, we further examined their effects on burst features. CNQX treated control neural networks decreased the number of bursts per well, while this was unchanged in SLC35A2^S304P/Y^ and SLC35A2^-/Y^ networks (Supplementary Fig. 10). There was also an observed increase in mean burst duration, inter-spike intervals (within bursts), and a decrease in the average spike frequency (within bursts) in CNQX treated control networks (Supplementary Fig. 10). In contrast, CNQX treated SLC35A2^S304P/Y^ and SLC35A2^-/Y^ networks did not affect mean burst duration or average spike frequency (within bursts), whereas it very slightly increased mean-interspike interval (within bursts) only in SLC35A2^-/Y^ (Supplementary Fig. 10). Bicuculline had marginal effects on burst dynamics across all networks. As expected, treatment of neural networks with TTX abolished all spontaneous activity (Supplementary Fig. 10 and Fig. 11). A summary of the effects of CNQX, bicuculline and TTX on spike and burst features are found in Supplementary Fig. 10-12.

Taken together, these data suggest that control and *SLC35A2* LOF neural networks *in vitro* contain both excitatory and inhibitory cell types. Furthermore, networks treated with CNQX and bicuculline suggest that excitatory neurons mediate MFR and bursting phenotypes where both excitatory and inhibitory neuron activity mediate network synchrony. Importantly, these data reveal a possible inhibitory/excitatory imbalance in SLC35A2^S304P/Y^ and SLC35A2^-/Y^ variant networks compared to the control.

### Loss of SLC35A2 skews differentiation towards the GABAergic fate

Longitudinal MEA data and pharmacologic perturbation of neural networks reveal distinct firing activity and patterns between control and both SLC35A2^S304P/Y^ and SLC35A2^-/Y^ networks. The decreased activity and asynchronous firing observed in SLC35A2^S304P/Y^ and SLC35A2^-/Y^ networks in addition to differential response to pharmacologic agents suggests an inhibitory/excitatory imbalance. To characterize early developmental trajectories (day 12 and 20) and neural composition in mature neural networks (day 20 and 84) we assessed the expression of various pallial, subpallial, glutamatergic and GABAergic specific markers (Fig. 6).

**Figure 6.**
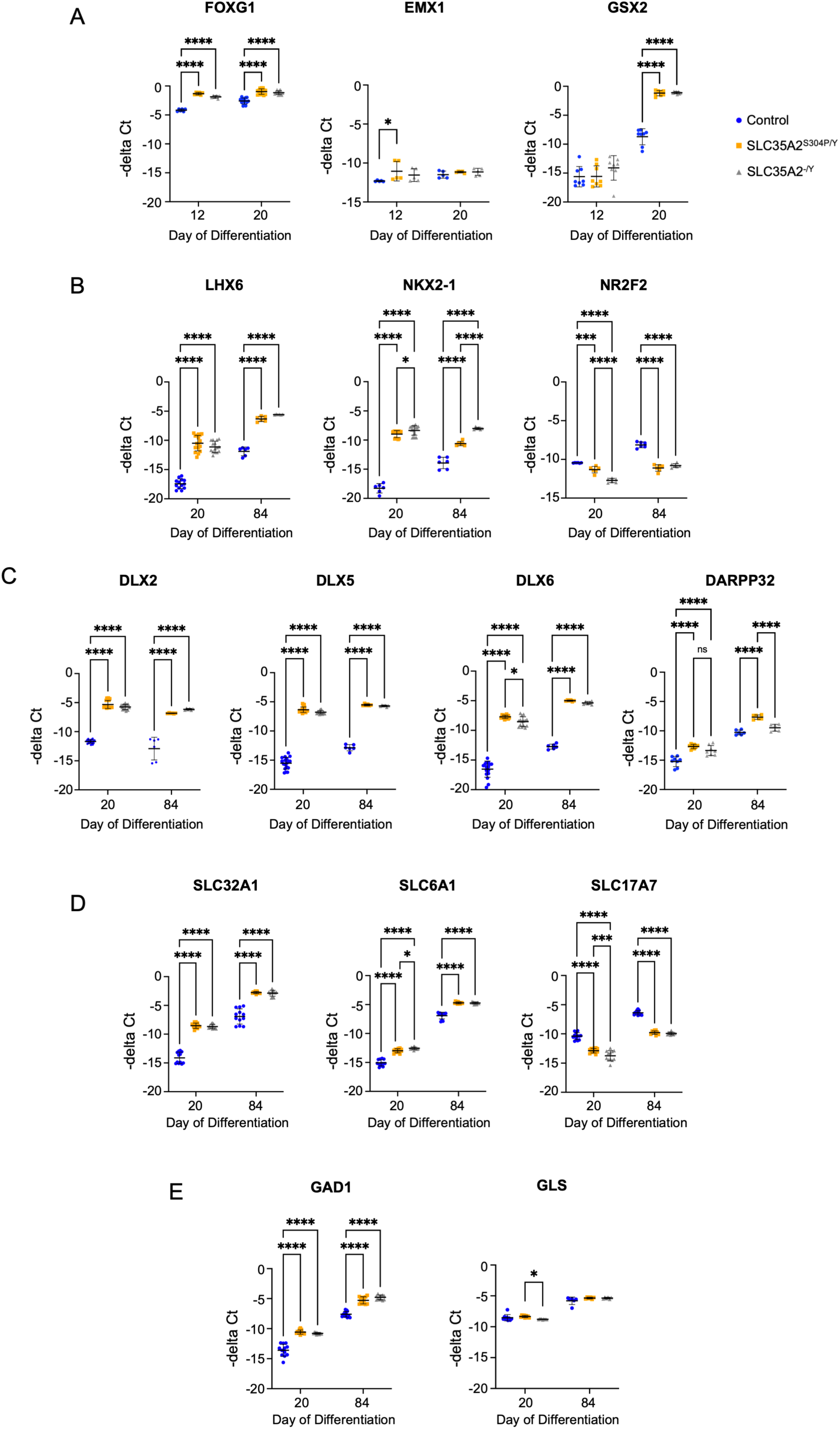
Characterization of early neurodevelopmental trajectories and neural composition. **A-E)** Quantification of mRNA expression levels representative of telencephalic development, pallium/subpallium, and GABAergic/glutamatergic specific markers using qRT-PCR. Data is representative of at minimum two pooled independent differentiations with each replicate run in triplicate per genotype. Each dot represents a single data point. Control is represented as blue circles, SLC35A2^S304P/Y^ as orange squares, and SLC35A2^-/Y^ as grey triangles. **(A)** Markers specifying the telencephalon (*FOXG1*), pallium (*EMX1*), and subpallium/lateral ganglionic eminence (LGE): *GSX2*. **(B)** Markers specific to the medial ganglionic eminence (MGE: *LHX6*, *NKX2-1*) and caudal ganglionic eminence (CGE: *NR2F2*). **(C)** GABAergic specific transcription factors important for differentiation and fate specification: *DLX2*, *DLX5*, *DLX6.* Medium spiny neuron marker: *DARPP32*. **(D)** Transporter specific markers. GABAergic: *SLC32A1*/VGAT (vesicular GABA transporter), *SLC6A1*/GAT1 (GABA transporter 1); Glutamatergic: *SLC17A7*/VGLUT1 (vesicular glutamate transporter 1). **(E)** Enzymatic markers for neurotransmitter synthesis. GABAergic: *GAD1* (glutamate decarboxylase 1). Glutamatergic: *GLS* (glutaminase). Statistics: Two-way ANOVA with Tukey’s to correct for multiple comparisons. * = P≤0.05; ** = P≤0.01; *** = P≤0.001; **** = P≤0.0001.

**Figure 7.**
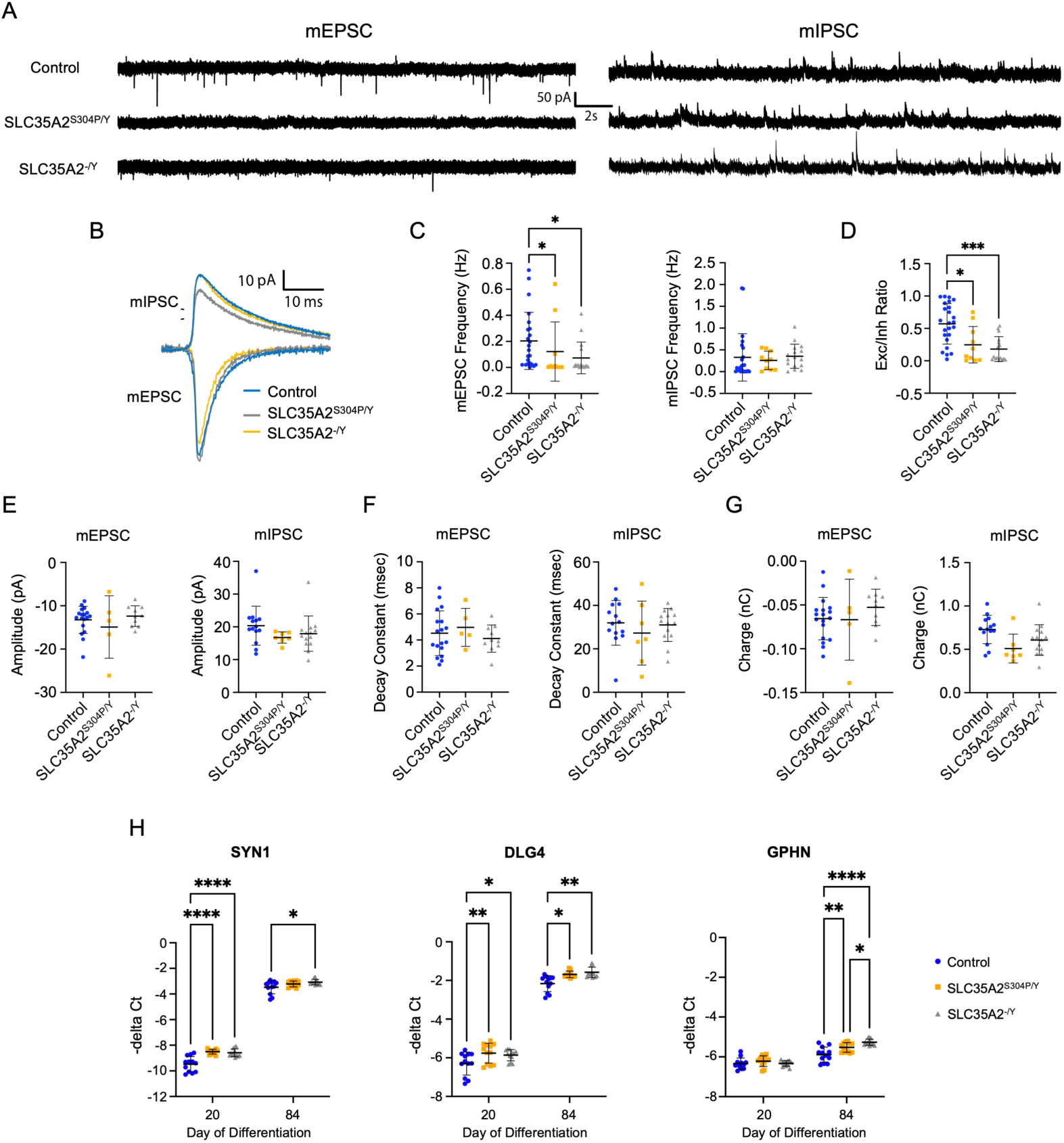
Whole-cell recordings reveal an enrichment in inhibitory synapses in SLC35A2^S304P/Y^ and SLC35A2^-/Y^ neurons. **(A)** Representative traces of miniature excitatory post-synaptic currents (mEPSC; left) and miniature inhibitory post-synaptic currents (mIPSC; right) recorded from the same neuron by holding voltage at −60 mV and 0 mV, respectively. Five-minute-long recordings were performed in whole-cell voltage-clamp mode in the presence of 100 nM TTX to eliminate neurotransmitter release caused by action potential-dependent network activity. **(B)** Average mEPSC and mIPSCs obtained from control (blue), SLC35A2^S304P/Y^ (grey), and SLC35A2^-/Y^ (yellow) neurons. **(C)** Event frequencies of mEPSC and mIPSCs (Hz). **(D)** Proportion of mEPSC to mIPSC events recorded per neuron for each genotype. Statistics for C and D: Kruskal-Wallis test followed by Dunn’s test to correct for multiple comparisons. **(E)** Amplitude of mEPSC and mIPSC events (pA). **(F)** Decay constant for mEPSC and mIPSC (msec). **(G)** Average charge transfer for mEPSC and mIPSC (nC). Synaptic data is representative of two pooled independent differentiations with each dot representing recordings from a single neuron. Statistics for E through G: One-way ANOVA with Tukey’s to correct for multiple comparisons. **(H)** Expression of synaptic markers. Pan-synaptic: *SYN1*; Excitatory post-synaptic: *DLG4*; Inhibitory post-synaptic: *GPHN*. Data is representative of two pooled independent differentiations each containing a technical replicate. Each replicate was run in triplicate per genotype. Each dot represents a single data point. Statistics for H: Two-way ANOVA with Tukey’s to correct for multiple comparisons. * = P≤0.05; ** = P≤0.01; *** = P≤0.001; **** = P≤0.0001.

During early neurodevelopment, all networks highly express *FOXG1*, a marker for the developing telencephalon (Fig. 6A). No difference in *EMX1* (pallial) expression is detected between the cultures, whereas *SLC35A2* LOF variant networks demonstrate a significant increase in *GSX2* expression - a general subpallial marker that also specifies the lateral ganglionic eminence (LGE) (Fig. 6A). Additional markers of the medial ganglionic eminence (MGE; *LHX6*, *NKX2-1*) and the caudal ganglionic eminence (CGE; *NR2F2*) reveal increased expression levels of *NR2F2* in control neurons, whereas SLC35A2^S304P/Y^ and SLC35A2^-/Y^ neurons express higher levels of *NKX2-1* and *LHX6* (Fig. 6B). Transcription factors that regulate the differentiation and fate specification of GABAergic neurons (*DLX2, DLX5, DLX6*) were also upregulated in SLC35A2^S304P/Y^ and SLC35A2^-/Y^ neural cultures compared to the control suggesting an enriched proportion of inhibitory neurons in *SLC35A2* LOF networks (Fig. 6C). Interestingly, SLC35A2^S304P/Y^ neural cultures expressed high levels of *DARPP32*, a marker for medium spiny neurons (GABAergic neurons of the striatum) which was not observed in SLC35A2^-/Y^ (Fig. 6C). This suggests a possible divergence in pathway regulation between SLC35A2^S304P/Y^ and SLC35A2^-/Y^ variants.

We next assessed functional markers of inhibitory and excitatory neurons. The mRNA expression levels of pan-GABAergic specific markers [vesicular GABA transporter (*SLC32A1*, VGAT), GABA transporter (*SLC6A1*, GAT1), and glutamate decarboxylase (*GAD1*; GAD67)] were all upregulated in SLC35A2^S304P/Y^ and SLC35A2^-/Y^ neural cultures compared to control (Fig. 6, D and E). Early differences are detected at day 20 and persist in mature neurons at day 84. The pan-glutamatergic marker *SLC17A7*, which encodes the vesicular glutamatergic transporter (vGLUT1), is downregulated in SLC35A2^S304P/Y^ and SLC35A2^-/Y^ cultures, whereas no discernible difference is detected in the expression of glutaminase (*GLS*), a glutamate synthesizing enzyme (Fig. 6, D and E).

To confirm mRNA expression levels reflects protein levels, we quantified the proportion of GABAergic neurons in day 52 and 86 cultures (Supplementary Fig. 13). At day 52, 9% of control, 31% of SLC35A2^S304P/Y^, and 36% of SLC35A2^-/Y^ neurons stained positively for GABA. This increased significantly to 68% of control, 86% of SLC35A2^S304P/Y^, and 91% of SLC35A2^-/Y^ neurons at day 86. The higher relative proportion of GABAergic neurons in *SLC35A2* LOF neural networks quantified at mRNA levels and validated with immunofluorescent staining suggests SLC35A2 may regulate cell fate specification. We also confirmed that neural networks all express markers of deep (TBR1, CTIP2) and upper layer (BRN2) cortical markers (Supplementary Fig. 14 and 15).

### *SLC35A2* LOF network phenotypes are driven by an enrichment in inhibitory synapses

While we observe an enriched inhibitory population in the *SLC35A2* LOF networks, intrinsic excitability and/or synaptic differences could also contribute to the observed network phenotypes. There were minor differences in the membrane properties between genotypes, but no differences in passive or active properties expected to change excitability were observed (Supplementary Fig. 16). Therefore, we next explored the role of synaptic changes by recording miniature synaptic currents (Fig. 7, A and B). Miniature excitatory post-synaptic currents (mEPSC) and miniature inhibitory post-synaptic currents (mIPSC) were recorded sequentially from the same neuron in the presence of TTX, a voltage-gated sodium channel blocker. TTX eliminates neurotransmitter release caused by action potential-dependent network activity and can identify changes occurring specifically at the synapse, which can be mediated by pre- and/or post-synaptic alterations. We investigated the involvement of both by measuring the frequency and amplitude of mEPSC and mIPSC. Changes in the frequency of miniature synaptic currents represent a relative change in the number of synaptic vesicles released at the pre-synaptic membrane whereas changes in amplitude suggest a relative change in the number of receptors on the post-synaptic membrane. An increase in mEPSC frequencies were detected in the isogenic control neurons whereas no difference in mIPSC frequencies were detected between the genotypes (Fig. 7C). However, when considering the total number of events, mEPSCs are enriched in control neurons whereas mIPSCs are enriched in SLC35A2^S304P/Y^ and SLC35A2^-/Y^ neurons (Fig. 7D). No differences in mEPSC or mIPSC amplitude, decay rate, or charge constant were observed (Fig. 7, E-G). These data provide evidence that neural network phenotypes are driven in part by changes in the number of excitatory and inhibitory vesicles released from the neuron termini in the control and *SLC35A2* LOF neural networks, respectively.

Although we did not detect any post-synaptic changes at the receptor level in the synaptic current recordings, we did detect differences in mRNA expression levels of post-synaptic density markers. We observed an increase in excitatory (*DLG4*) and inhibitory (*GPHN*) specific post-synaptic markers in *SLC35A2* LOF neural networks compared to the control (Fig. 7H). This suggests additional changes may occur post-synaptically related to the stabilization of glutamate receptors, strength of neurotransmission, synaptic organization, or plasticity. We also observed a small increase in the pan pre-synaptic marker Synapsin1 (*SYN1*) in SLC35A2^-/Y^ neurons compared to the control (Fig. 7H).

## Discussion

In this study, we provide a first in-depth assessment of the functional consequences of a patient identified missense variant in *SLC35A2* (SLC35A2^S304P/Y^) and knockout of *SLC35A2* (SLC35A2^-/Y^) using a human isogenic iPSC-derived neuron model. We integrated glycobiology, cellular and molecular analysis, network electrophysiology, and single cell electrophysiology for comprehensive *in vitro* phenotypic analyses. Through this multidisciplinary approach, we provide compelling evidence demonstrating that LOF variants in *SLC35A2* perturb neurodevelopment and neuronal network activity driven by preferential differentiation towards the GABAergic fate and associated pre-synaptic changes. These findings provide significant groundwork for further investigation of specific mechanisms underlying *SLC35A2* pathogenesis, paving the way for the development of targeted therapeutics.

MCDs are acquired during neurodevelopment. *SLC35A2* variants likely arise during this critical time that can drive underlying pathology. Our data reveal that *SLC35A2* LOF variant harboring cultures led to accelerated neurogenesis compared to the isogenic control as evidenced by the detection of premature, dysmorphic neural rosettes and ß3-tubulin positive neurons (Fig. 3, A and B). Given the ubiquity of glycosylation and the remarkable differences in complexity of N-glycan spectra in iPSCs and neurons (Fig. 1D and 2B), we speculate that aberrant glycosylation contributes to the accelerated neurogenic phenotype. In support of this hypothesis, it has been shown that the N-glycome in adult brain tissue is enriched for less complex, truncated N-glycans relative to human prenatal brain tissue,^37^ which is comparable to our observation in *SLC35A2* LOF variant cultures. Another study demonstrated that loss of N-glycan branching accelerates neurogenesis in mouse neural stem and progenitor cells.^38^ Further in-depth analysis demonstrating that the accelerated neurogenic phenotype observed in *SLC35A2* LOF variant cultures is a direct cause of the reduced complexity of N-glycans would be needed to confirm this relationship. Importantly, the reduced galactosylation phenotype in our iPSC-derived neuron model of *SLC35A2* variants aligns with similar findings in resected brain tissue from mild MCD patients, whole serum of individuals with CDG, and in *SLC35A2*-deficient cell lines ^2,24,33^ – regardless of the method utilized for glycan analysis (e.g., LC-MS, HPLC, lectin blots) – providing strong evidence that our *in vitro* model recapitulates features of human disease.

Differential glycosylation modifications have also been shown to modulate cell fate specification.^38,39^ Although the directed differentiation protocol we used results in a culture enriched with glutamatergic neurons,^28,31^ we observed a higher proportion of GABAergic neurons in *SLC35A2* LOF networks (Supplementary Fig. 13). It is possible that intrinsic differences in Wnt and/or SHH signaling, specifically low endogenous Wnt signaling or high SHH, is driving the ventralized fate in *SLC35A2* LOF variant networks.^40^ This enriched GABAergic population is likely contributing to the hypoactive, asynchronous network phenotype observed by *SLC35A2* LOF variants (Fig. 4, B and C), which may provide favorable conditions that could precipitate conditions for seizure initiation as seen in other epilepsy models.^41,42^ This hypothesis is in contrast to the conventional paradigm that epileptic seizures are a result of hyperactive, hypersynchronous events, however, inhibitory neuron activity and asynchronous networks have both been captured at seizure onset and during the ictal period.^43–51^ Kooshkhoo *et al*.^52^ investigated the differential effects of inhibitory neuron subtypes on seizure initiation and duration, elucidating VIP neurons as a possible target to reduce seizure duration. Enriched proportions and/or activity of inhibitory neurons have also been previously identified as the cellular and network underpinnings in other genetic models of epilepsy, Rett Syndrome, PACS1 syndrome, and autism spectrum disorders, further highlighting the alterations in number and activity of GABAergic neurons in mediating a spectrum of neurodevelopmental phenotypes.^53–59^

While we describe the role of inhibitory neurons to the epilepsy phenotype, other mechanisms may also contribute. Recent studies used *in vitro* electroporation to selectively knockout *SLC35A2* in embryonic mice targeting layer II/III cortical neurons.^60,61^ Since only a proportion of neurons would be targeted, this method captures the somatic mosaicism observed in affected individuals with an *SLC35A2* variant. Post-natal analyses reveal *SLC35A2* knockout neurons display cortical dyslamination with heterotopic neurons mispositioned in deep cortical layers and white matter, suggesting dysregulated radial migration of neural progenitors. Cortical dyslamination is a defining histopathological feature of MCDs, specifically FCDI and FCDII.^12^ Therefore, these data provide evidence that LOF mutations in *SLC35A2* disrupt neuron migration leading to cortical dyslamination consistent with the MCD phenotype, and possibly contributing to the seizure phenotype.

In addition to activity and synchrony metrics, our MEA analyses revealed differences in burst dynamics. Specifically, prolonged burst duration and increased inter-spike interval were detected in *SLC35A2* LOF networks compared to control (Fig. 4, D and E). These differences in burst features are comparable to those observed in a recent study modeling PACS1 syndrome, ^58^ where prolonged network bursts and increased interspike intervals (within bursts) were observed in PACS1 mutant neural networks compared to the control. This phenotype is reminiscent of differences in fast after hyperpolarization (fAHP) kinetics mediated by calcium- and voltage-dependent potassium channels, known as BK channels.^62,63^ Disrupted N-glycosylation of BK channels reduces their expression, localization and function.^64^ Therefore, it is possible that the reduced glycosylation detected in SLC35A2^S304P/Y^ and SLC35A2^-/Y^ neural networks may affect BK channel function leading to differential fAHP kinetics and altered burst dynamics.

The pathogenic variants in *SLC35A2* identified to date, collectively explain the greatest fraction of monogenic neocortical epilepsies, making it an ideal gene to advance the precision medicine paradigm. Pathogenic variants in *SLC35A2* include missense, frameshift indels, nonsense, and splice site mutations that are absent from control databases (EXAC, gnoMAD).^23,25,65^ All are theorized to cause epilepsy through LOF mechanisms impacting transporter expression, stability, localization, and/or function.^23,25,65^ Since SLC35A2 is the only known transporter of UDP-galactose from the cytosol into the Golgi where galactosylated glycans form, these disease-causing variants generate reduced galactosylated glycans (Fig. 1D and 2B).^2,65–67^ Galactose supplementation is a therapeutic option to potentially overcome the *SLC35A2* variant-induced transporter deficiency for some missense variants. Dietary D-galactose is currently used to treat phosphoglucomutase 1 deficiency (PGM1-CDG) underscoring its possible use for *SLC35A2*-associated epilepsy and/or *SLC35A2*-CDG. Clinical studies have investigated the efficacy of dietary galactose in eleven *SLC35A2*-CDG patients, five of which experience seizures, and in six cases of MOGHE.^68–70^ Based on these studies, there appears to be some benefit of galactose supplementation on seizure outcomes in approximately half of the patients, however, the sample size is too small to be able to draw any significant conclusions and the authors did not report a genotype relationship associated with treatment outcome. Nonetheless, we do hypothesize that galactose efficacy will be largely dictated by transporter localization to the Golgi and the nature of the mutation; the latter of which can greatly impact transporter kinetics and substrate recognition.^2,71^ Since the variants generated in this study (SLC35A2^S304P/Y^ and SLC35A2^-/Y^) do not encode SLC35A2 protein (Fig. 1B and C) they are likely unable to transport any UDP-galactose across the Golgi membrane and thus, galactose supplementation will have a negligible effect, if any. Further experimentation with protein coding, Golgi-localized missense variants would be more suitable to assess the effects of galactose supplementation in reversing the observed *SLC35A2*-associated phenotypes.

*SLC35A2* expression and Golgi localization are important indicators for predicting galactose response. In cases where SLC35A2 is not expressed or is mis-localized, gene-targeted therapies could be beneficial. The X-linked mode of inheritance of *SLC35A2* does complicate possible therapeutic strategies. Additionally, since all variants identified are somatic, only a fraction of cells will harbor the mutation. Previous work has identified that *SLC35A2* variants associated with MOGHE are present in both neurons and oligodendrocytes, with a particular enrichment in the latter, suggesting that the variant arose in neuroglial progenitors.^27^ However, the specific neuronal subtype has yet to be identified in human postmortem tissue. Our data suggest that GABAergic neurons may contribute to *SLC35A2*-mediated epilepsy. Lastly, *SLC35A2* variant enrichment in oligodendrocytes and the presence of an oligodendroglial-associated neuropathology (MOGHE) in brain tissue underscores their contribution in the pathogenesis of MCD, MOGHE and epilepsy associated with CDG cases. Future work on the functional characterization of variant harboring oligodendrocytes will be essential to our understanding of cell type specific effects of *SLC35A2* variants on neurodevelopmental and network phenotypes.

This is the first study to investigate the functional effects of patient-identified *SLC35A2* LOF variants using an isogenic human iPSC-derived neuron model. We find that *SLC35A2* LOF variants disrupt glycosylation, promote precocious neurodevelopment, skew neuronal fate, and generate hypoactive, asynchronous networks that may be in part driven by changes at the pre-synaptic terminal. Beyond its role as a UDP-galactose transporter, we identified a novel role for *SLC35A2* in regulating neurodevelopment and cell fate determination likely mediated by alterations in cellular glycosylation.

## Abbreviations

CDG: congenital disorder of glycosylation
CGE: caudal ganglionic eminence
CNQX: cyanquixaline
EOEE: early onset epileptic encephalopathy
fAHP: fast afterhyperpolarization
FCDI: focal cortical dysplasia type I
FCDII: focal cortical dysplasia type II
HMEG: hemimegalencephaly
iPSC: induced pluripotent stem cell
LOF: loss of function
MCD: malformation of cortical development
mMCD: mild malformation of cortical development
MEA: multielectrode array
MGE: medial ganglionic eminence
mEPSC: miniature excitatory postsynaptic current
MFR: mean firing rate
mIPSC: miniature inhibitory postsynaptic current
MOGHE: malformation of cortical development with oligodendroglial hyperplasia in epilepsy
ProA: procainamide hydrochloride
STTC: spike train tiling coefficient
TTX: tetrodotoxin
UDP: uridine diphosphate
VAF: variant allele fraction

## Acknowledgements

We thank Jeanne Loring for providing the control human iPSC line. We thank the Columbia Stem Cell Core for their assistance with generating the CRISPR edited iPSCs. We also like to thank the UNC Neuroscience Microscopy Core (RRID:SCR_019060), supported, in part, by funding from the NIH-NINDS Neuroscience Center Support Grant P30 NS045892 and the NIH-NICHD Intellectual and Developmental Disabilities Research Center Support Grant 50 HD103573.

## Supplementary material

Supplementary material is available for download.

