## SupplementaryMaterial for "SLC35A2 loss of function variants affect glycomic signatures, neuronal fate, and network dynamics"

#### Supplementary Methods

*CRISPR-Cas9 genome editing*
The 713-5 isogenic control was edited to harbor a patient-specific missense variant in *SLC35A2* as identified in our previous study [c.910T>C p.(Ser304Pro)].[1] Guide RNAs (gRNA) were designed using the algorithms CHOPCHOP and CRISPRscan to minimize off-target effects.[2-4] The guide RNA (GGTGGACAGCACAATGGACA) was synthesized within a genomic block (gBlock) under the U6 promoter and cloned into the pCR-BluntII-TOPO vector. gBlock sequence and guide RNA (underlined), as follows:
TGTACAAAAAAGCAGGCTTTAAAGGAACCAATTCAGTCGACTGGATCCGGTACCAAG
GTCGGGCAGGAAGAGGGCCTATTTCCCATGATTCCTTCATATTTGCATATACGATACAA
GGCTGTTAGAGAGATAATTAGAATTAATTTGACTGTAAACACAAAGATATTAGTACAA
AATACGTGACGTAGAAAGTAATAATTTCTTGGGTAGTTTGCAGTTTTAAAATTATGTTT
TAAAATGGACTATCATATGCTTACCGTAACTTGAAAGTATTTCGATTTCTTGGCTTTATA
TATCTTGTGGAAAGGACGAAACACCGGTGGACAGCACAATGGACAGTTTTAGAGCTA
GAAATAGCAAGTTAAAATAAGGCTAGTCCGTTATCAACTTGAAAAAGTGGCACCGAG
TCGGTGCTTTTTTTCTAGACCCAGCTTTCTTGTACAAAGTTGGCATTA.
The gRNA, repair template (GTCTGGGGCGTGGTGCTCAACCAGGCCTTCGGCGGGCTA
CTGGTGGCTGTGGTTGTCAAGTACGCTGACAATATCCTCAAGGGCTTTGCCACCTCCC
TG**C**CCATTGTGCTGT**CAA**CTGTTGCCTCCATTCGCCTCTTTGGCTTCCACGTGGACCC
ATTATTTGCCCTTGGCGCTGGACTCGTCATTGGTGCTGTCTACC), and AltR modified Cas9 RNP (Alt-R *S.p.* HiFi Cas9 Nuclease V3, Integrated DNA Technologies) were electroporated into hiPSC using the Neon Transfection System (Thermo Scientific). Transfected cells were selected with puromycin and sorted into single cells. Resulting colonies were lysed using the Kapa Express Extract Kit (Roche Diagnostics; 07961600001), amplified with MyTaq HS Mix (Meridian Bioscience; BIO25045), and purified with ExoSAP-IT PCR Cleanup Reagent (Applied Biosystems; 75001) prior Sanger sequencing to select variant positive clones. A knockout hiPSC line, harboring a two base pair deletion (c.911_912del), was also selected for functional studies.

*Glycan peak assignment*
Glycan HPLC peaks were assigned using a comparative approach with known standards. Specifically, we deglycosylated IgGs and RNase B followed by the release, purification, and labeling of N-glycans in the same manner as for those released from our sample. The specific N-glycans in our sample were then determined based on the elution time of the N-glycans released from the IgG and RNase B as described in the literature.[5] For calibration, to ensure consistent elution times and reliable peak identification across runs, IgG and RNase B glycans were always included along with our experimental samples.

*Mouse cortical astrocyte dissociation and culture*
Mouse cortical astrocytes were harvested from C57BL/6 P0 pups based on a previously published protocol with slight modifications.[6] In brief, the brain was isolated following decapitation of P0 pups and submerged in ice cold Hibernate-A (Gibco; A1247501). All extraneous structures were removed (e.g., olfactory bulbs, midbrain, hippocampus, meninges) leaving the cortices behind. Cortices were collected in a 15 mL conical tube containing ice-cold Hibernate-A and transferred into a 60 mm dish where they were finely minced into 1-2 mm pieces with a sterile blade. The mince was transferred to a 15 mL tube and allowed to settle. Excess media was removed and

cortical tissue was incubated with 2 mL of dissociation solution [20 units/mL of papain (Worthington Biochemical, LK003178) and 200 units/mL DNase (Worthington Biochemical, LK003172) reconstituted in Hibernate-A] for 20 minutes in a 37$^{o}$C waterbath with agitation every two minutes. The suspension was centrifuged at 0.3 rcf for 5 minutes and the pellet resuspended with 3 mL of Hibernate-A by vigorously pipetting 10x with a P1000 pipette. The suspension was incubated for three minutes to allow debris to settle. The supernantant (contains astrocytes) was then collected and transferred to a 15 mL tube. This collection step was repeated with another 3 mL of Hibernate-A and vigorously pipetted for a total of 6 mL supernatant. The suspension was then centrifuged at 0.3 rcf for 5 minutes to pellet the cells, resuspended in astrocyte media [DMEM (Gibco, 11995065), 10% FBS (Gibco, 10438026), 1x NEAA (Gibco, 11140050), 1% Pen/Strep (Gibco, 15140122)] and plated into a T75 flask. Astrocytes were expanded until passage 4 at which point they were treated with 10 uM of cytosine ß-D-arabinofuranoside hydrochloride (Ara-C; Sigma; C6645) prior to cryopreservation with freezing media (90% astrocyte media, 10% DMSO).

*Intrinsic excitability and action potential properties*

Membrane excitability was measured using current clamp recording. The pipette solution contained 130 mM $CH_3KO_3S$, 10 mM $CH_3NaO_3S$, 1 mM $CaCl_2$, 10 mM EGTA, 10 mM HEPES, 5 mM MgATP and 0.5 mM $Na_2GTP$ (pH 7.3, 305 mOsm). A -14 mV liquid junction potential between pipette and external solutions was calculated empirically, and the correction applied before the experiment. Resting membrane potential was measured immediately following establishment of the whole-cell configuration. During recordings, current (< 100 pA) was manually injected to hold the cells at approximately −60 mV. Membrane resistance and capacitance were calculated from the membrane potential changes in response to 1 s duration hyperpolarizing current steps that decreased in -5 pA increments. Action potentials were evoked and rheobase obtained using 1 s duration depolarizing current steps that increased incrementally by 5 pA. An action potential was defined as a transient depolarization of the membrane which had a minimum rise rate > 10 mV/ms and reached a peak amplitude > 0 mV. Action potential characteristics were measured from the first action potential at rheobase. The threshold potential was measured at the point where the voltage increases at a rate greater than 10 ms/mV. The duration was calculated from the full width at the half maximum voltage. For this calculation, the amplitude was measured from 0 mV to the peak potential. The maximum number of action potentials was measured from a 1 s current step.

Quantification was carried out using custom scripts written for Igor Pro (Wavemetrics, USA) and R v.3 (www.R-project.org). Statistical comparisons were made using the tests indicated in the Results section. P values < 0.05 were considered significant.

### Supplementary Figure 1

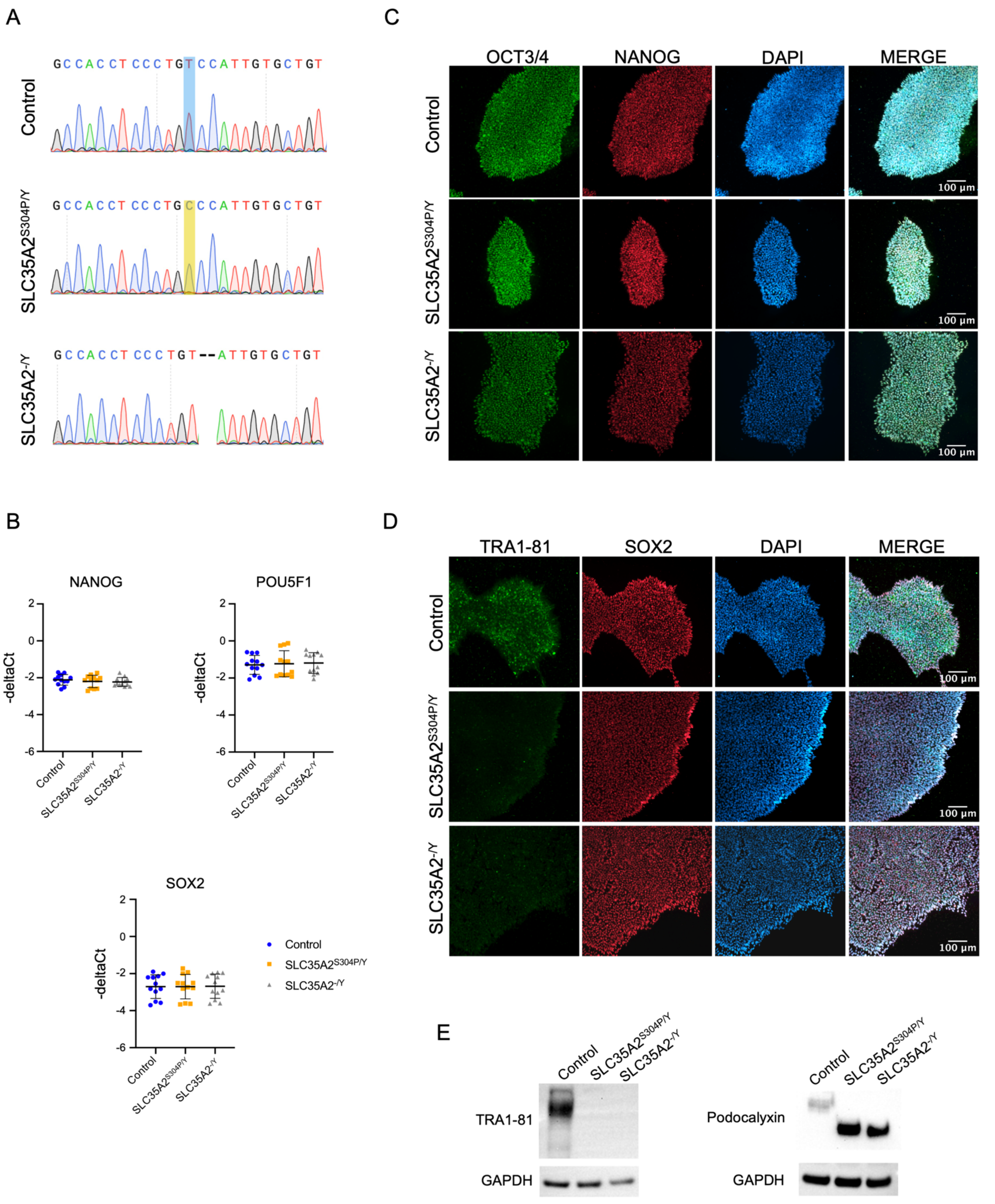

**Figure S1. Sequence validation of CRISPR/Cas9-edited iPSCs and characterization of pluripotency. (A)** Sanger sequencing of the variant site. Reference nucleotide is highlighted in blue (control), the patient identified missense variant is highlighted in yellow ($SLC35A2^{S304P/Y}$), and nucleotide deletions in the CRISPR-induced indel are represented by dashes ($SLC35A2^{-/Y}$). **(B)** mRNA expression levels of pluripotent markers. Data is representative of four pooled independent replicates. Each marker was run in triplicate per sample. Statistics: One-way ANOVA with Tukey's multiple comparisons. Data is represented as mean and standard deviation. Immunofluorescent staining for pluripotent markers **(C)** OCT3/4 and NANOG and **(D)** TRA-1-81 and SOX2. Nuclei were stained with DAPI. **(E)** Western blot. No observable detection of TRA-1-81 (left) in $SLC35A2^{S304P/Y}$ and $SLC35A2^{-/Y}$, despite detectable total amounts of podocalyxin (right). This suggests loss of the carbohydrate epitope (TRA-1-81) on the podocalyxin protein. GAPDH was used as a loading control. Un-cropped, full-length blots are provided in the Supplementary Materials.

#### Supplementary Figure 2

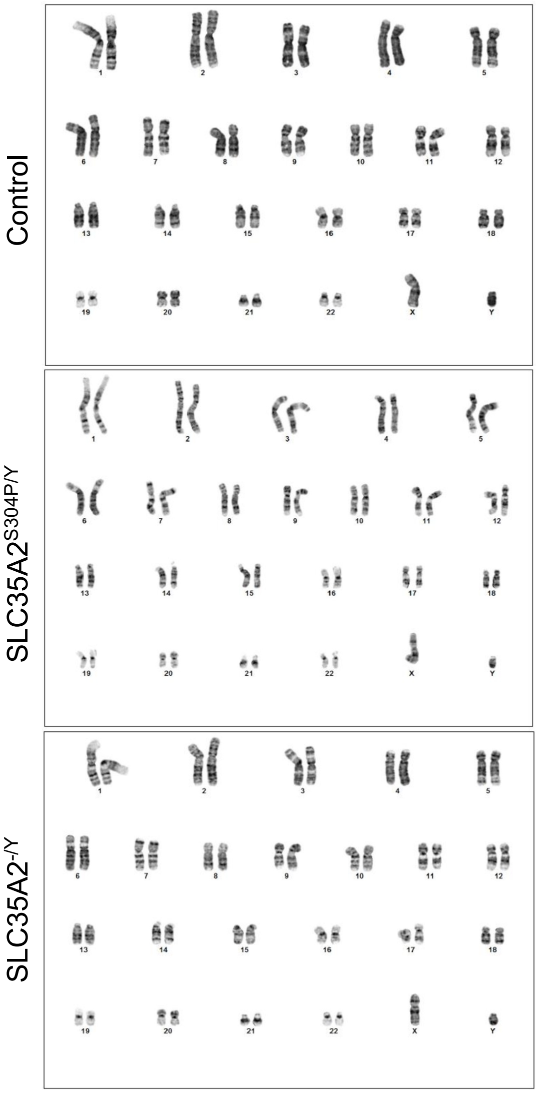


**Figure S2. Cytogenic analysis reveals normal karyotype.** iPSCs were subject to G-band metaphase analysis.

#### Supplementary Figure 3

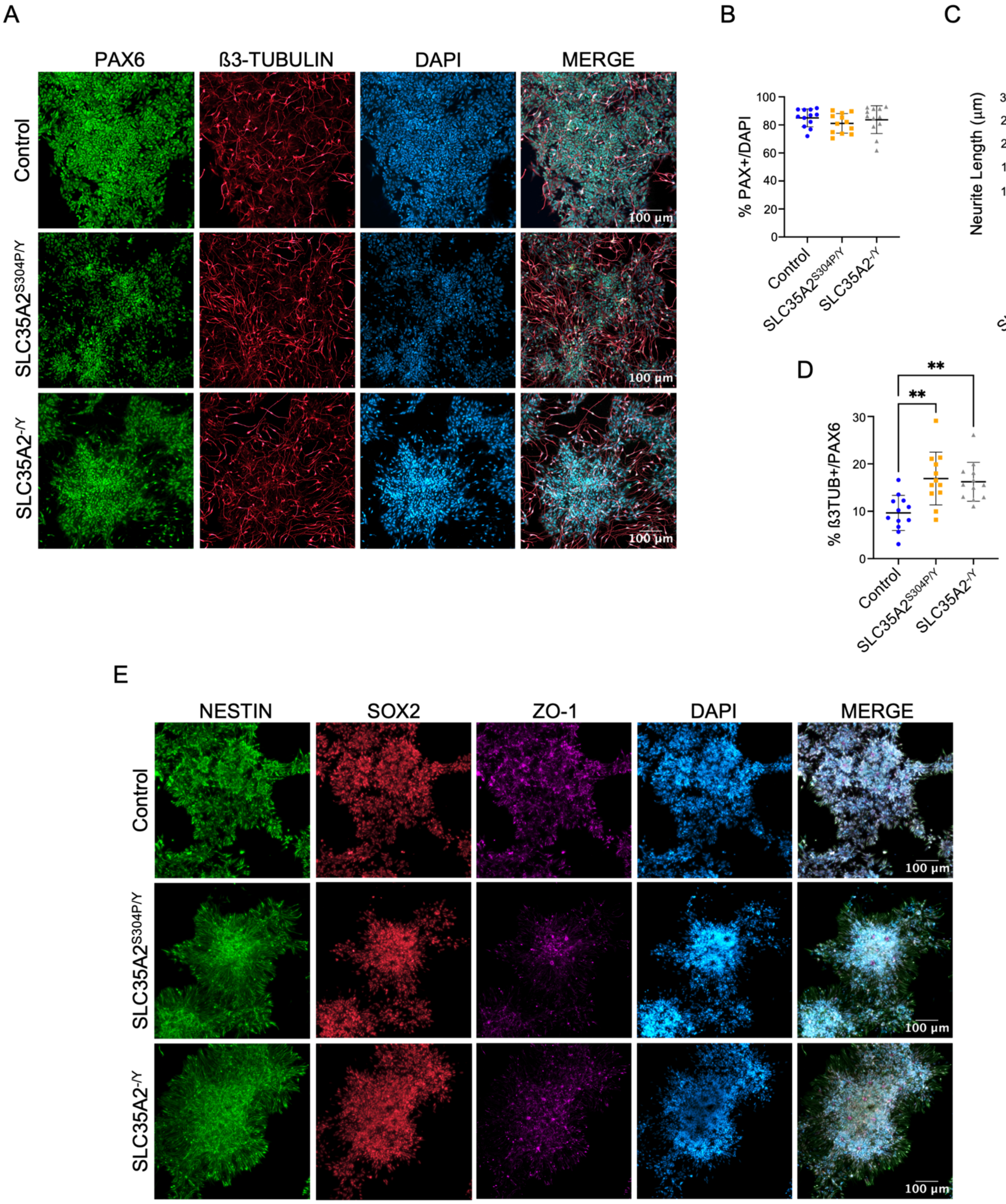

**Figure S3. Directed differentiation of iPSCs to cortical neurons. (A)** Day 20 cultures were stained for neural progenitors (PAX6), immature neurons (ß3-TUBULIN), and nuclei (DAPI). **(B)** Neural induction efficiency was quantified by the percent PAX6+ staining of total nuclei. **(C)** Quantification of neurite length at day 20 of differentiation (in μm). **(D)** Quantification of the proportion of ß3-TUBULIN+ neurons of total PAX6+ nuclei. **(E)** Day 25 cultures were stained for neural stem cell (SOX2, NESTIN), and neuroepilethelial (ZO-1) markers. Data from B and D are representative of four independent differentiations. Total nuclei counted: control n=17,592; $SLC35A2^{S304P/Y}$ n=12,001 $SLC35A2^{-/Y}$ n=13,330. Data from C is representative of three independent differentiations. Eight regions of interest (ROI) were captured per coverslip. Each dot represents quantification from one ROI. Statistics: One-way ANOVA with Tukey's multiple comparisons. Data is represented as mean and standard deviation. * = $P \leq 0.05$; ** = $P \leq 0.01$; *** = $P \leq 0.001$ **** = $P \leq 0.0001$.

#### Supplementary Figure 4

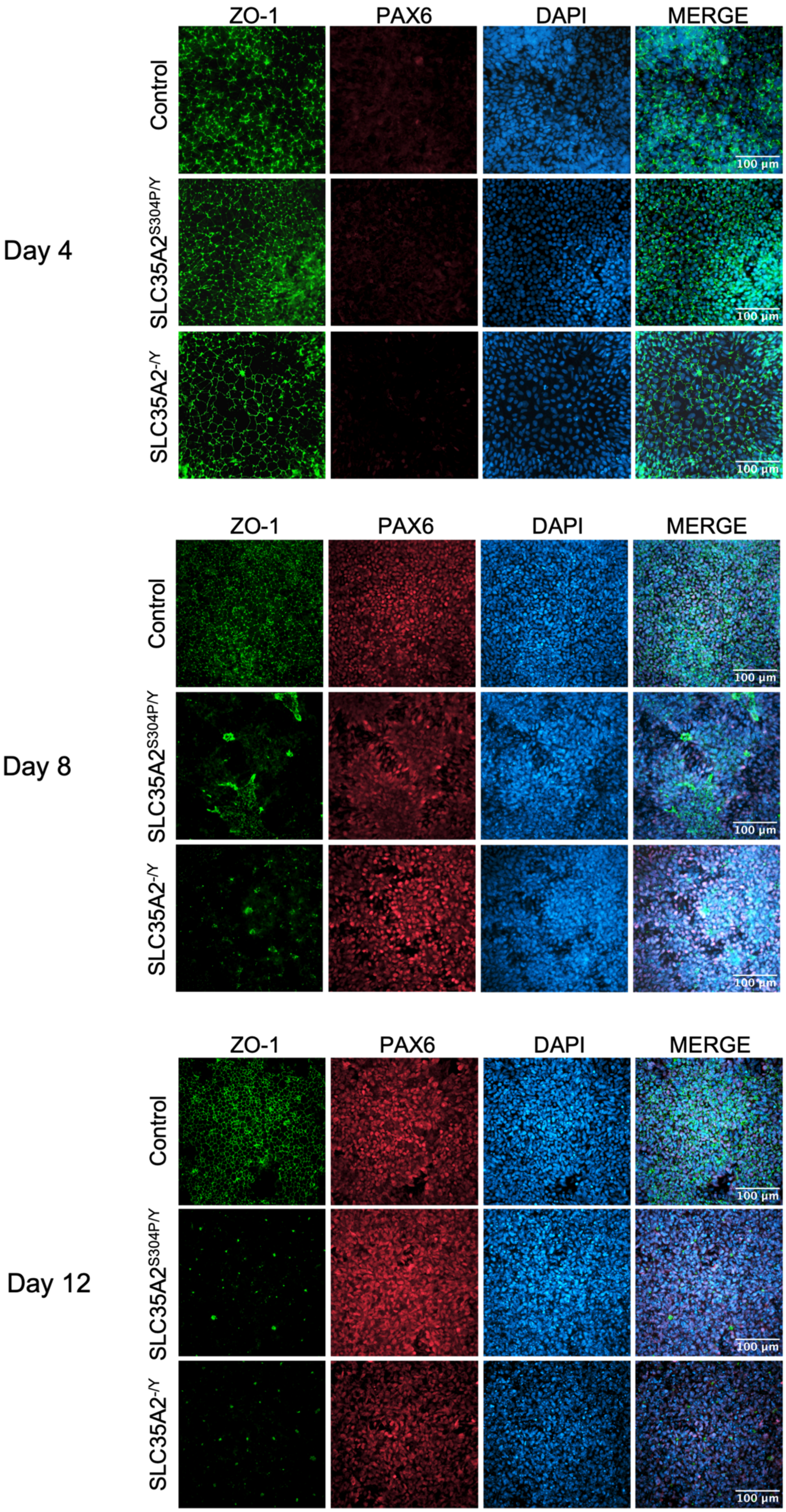

**Figure S4. Structural differences in neural rosettes from SLC35A2$^{S304P/Y}$ and SLC35A2$^{-/Y}$ variant harboring cultures.** Control, SLC35A2$^{S304P/Y}$ and SLC35A2$^{-/Y}$ cultures were fixed at day 4, 8, and 12 after neural induction. Lumen formation of neural rosettes among PAX6+ cells was detected by ZO-1 staining. Nuclei were detected with DAPI. Note the precocious lumen formation in the *SLC35A2* mutant cultures.

**Supplementary Figure 5**

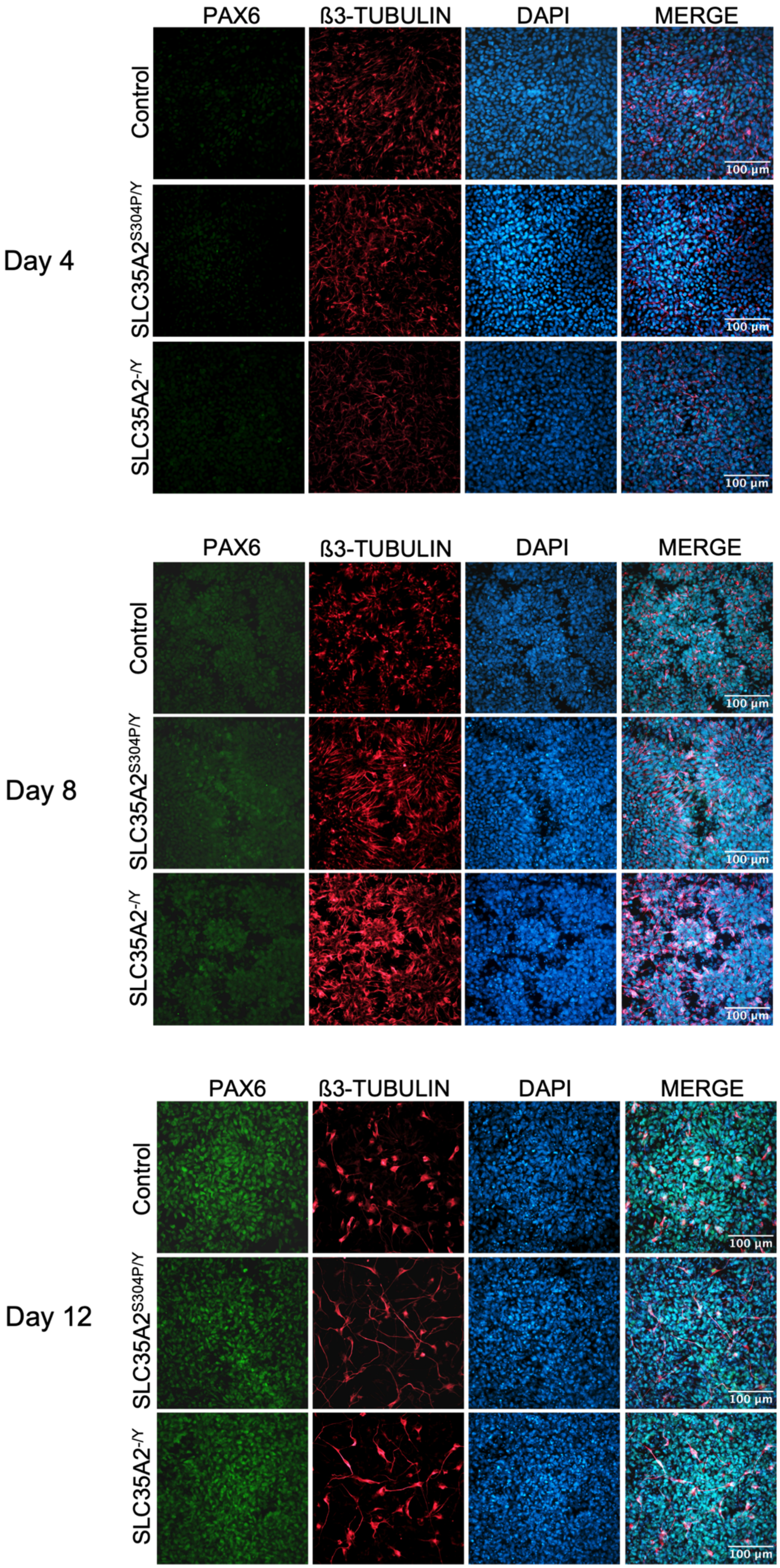

**Figure S5. Enhanced neurogenesis detected in SLC35A2$^{S304P/Y}$ and SLC35A2$^{-/Y}$ variant harboring cultures.** Control, SLC35A2$^{S304P/Y}$ and SLC35A2$^{-/Y}$ cultures were fixed at day 4, 8, and 12 after neural induction. Immature neurons were detected by staining for ß3-TUBULIN, neural progenitors were stained with PAX6 and nuclei were detected with DAPI.

#### Supplementary Figure 6

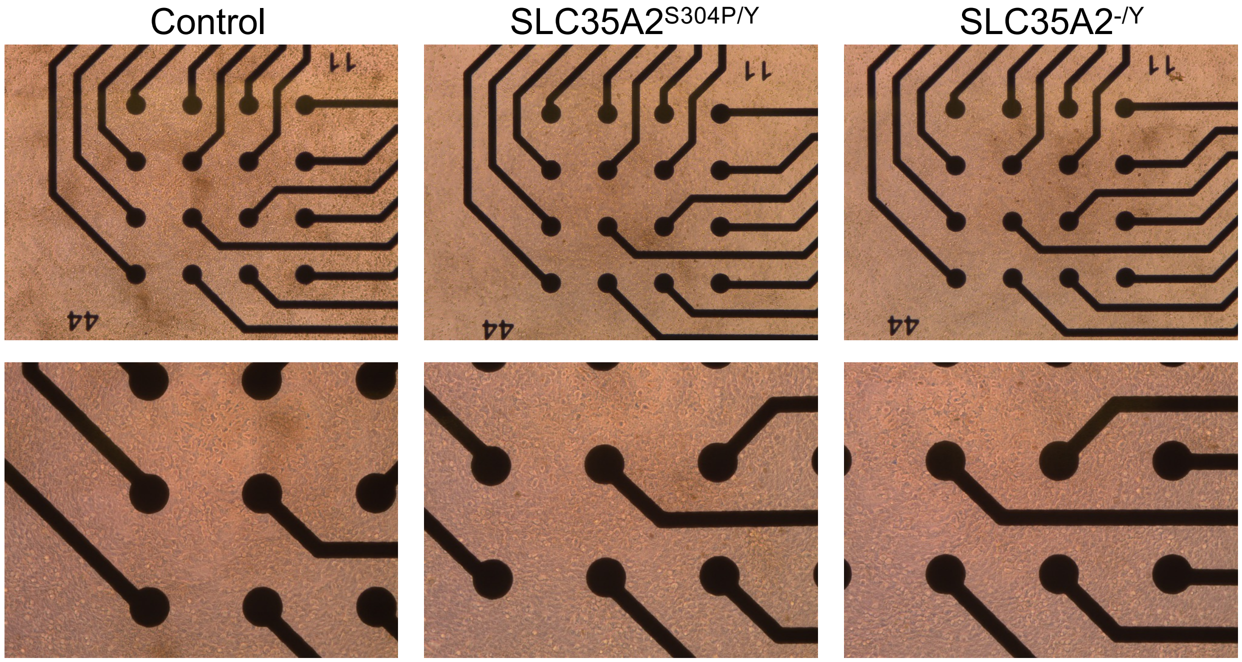


**Figure S6. Representative images of MEA wells.** Neurons were co-cultured with mouse astrocytes onto 48-well MEA plates containing 16 electrodes per well. Images were captured at day 84 of differentiation (or 60 days of co-culture conditions). Top row represents 5x magnification and bottom row represents 10x magnification.

#### Supplementary Figure 7

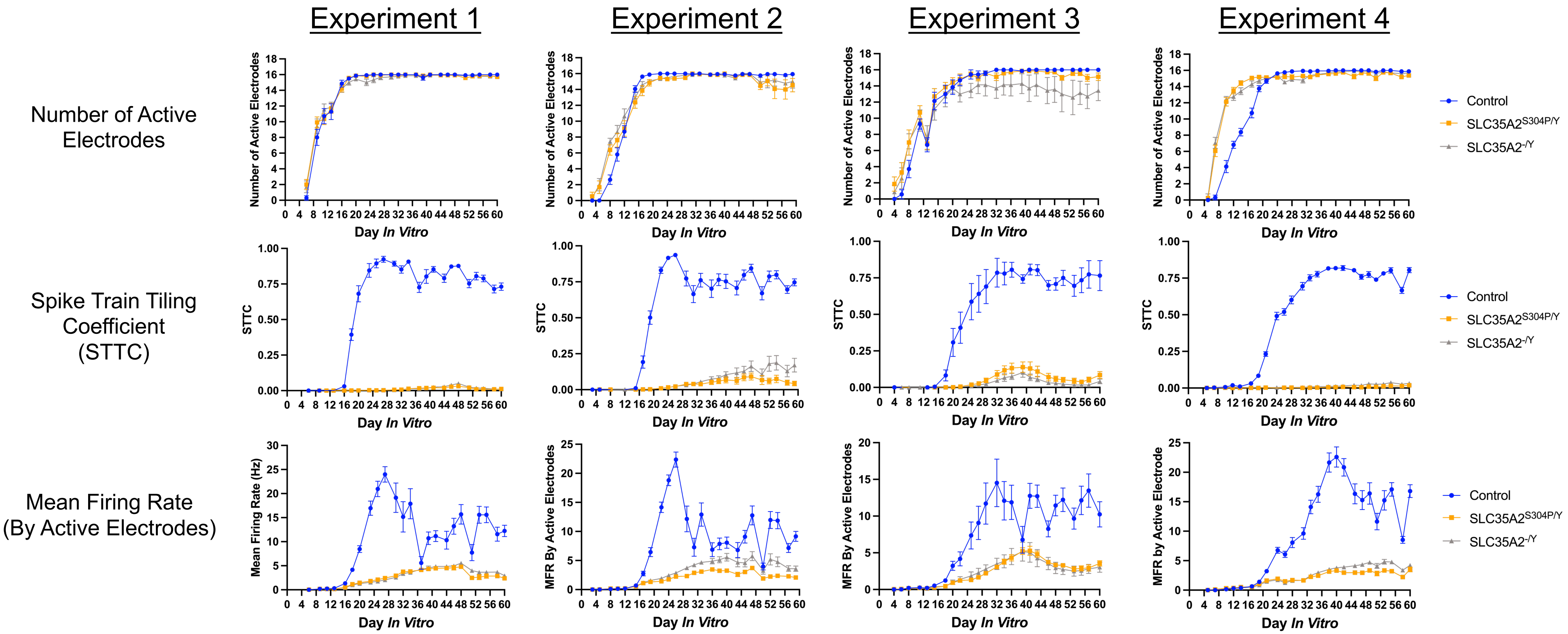


**Figure S7. Spike features extracted from each longitudinal MEA experiment reveal differences in neural activity and synchrony measures.** Each experiment represents the pooled average of an independent differentiation. For each feature, all wells per genotype were pooled, including replicates A and B, if present, for each MEA plate. Refer to Supplementary Table 5 for details regarding experimental replicates and number of wells plated per replicate on MEA. The mean and standard error are shown for each day *in vitro*, which represent the number of days following plating on MEA. Blue circles represent the control, orange squares represent SLC35A2$^{S304P/Y}$, and grey triangles represent SLC35A2$^{-/Y}$.

### Supplementary Figure 8

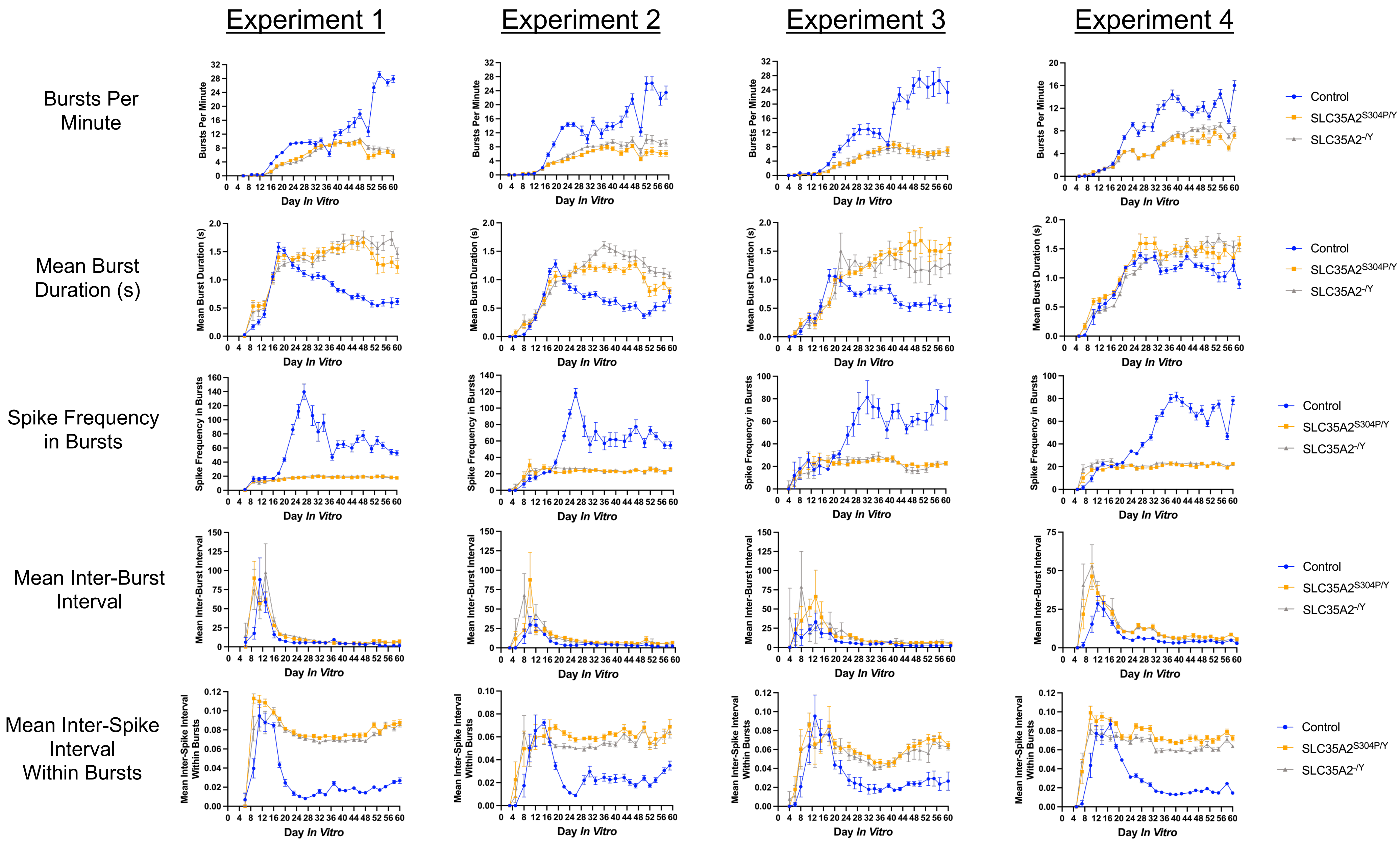

**Figure S8. Burst features extracted from each longitudinal MEA experiment reveal differences in firing dynamics**. Each experiment represents the pooled average of an independent differentiation. For each burst feature, all wells per genotype were pooled, including replicates A and B, if present, for each MEA plate. Refer to Supplementary Table 5 for details regarding experimental replicates and number of wells plated per replicate on MEA. The mean and standard error are shown for each day *in vitro*, which represent the number of days following plating on MEA. Blue circles represent the control, orange squares represent SLC35A2$^{S304P/Y}$, and grey triangles represent SLC35A2$^{-/Y}$.

#### Supplementary Figure 9

Experiment 1

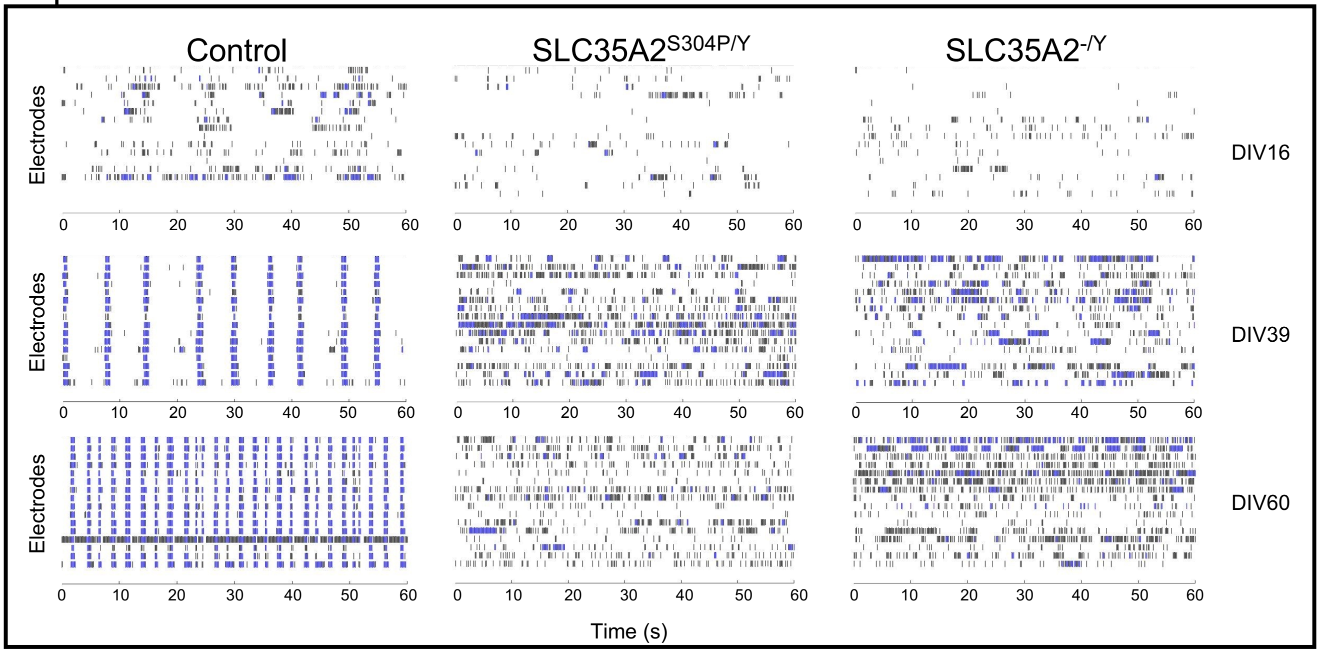


Experiment 2

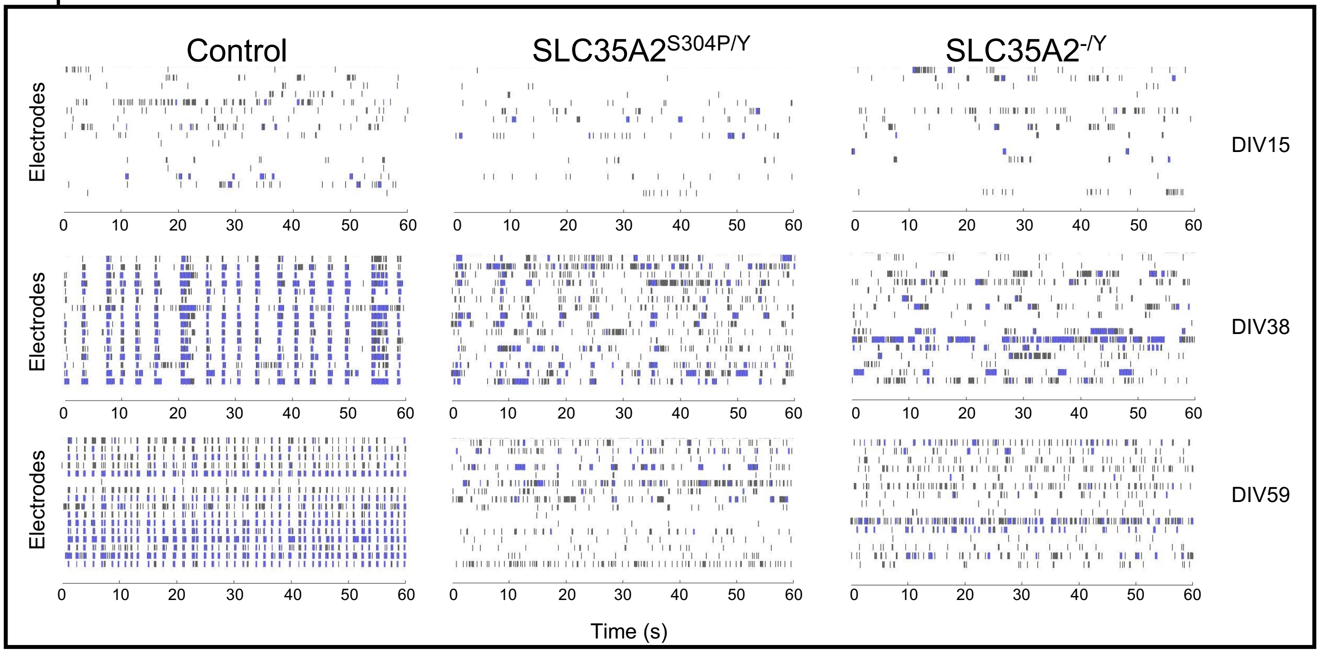

#### Supplementary Figure 9 (cont'd)

Experiment 3

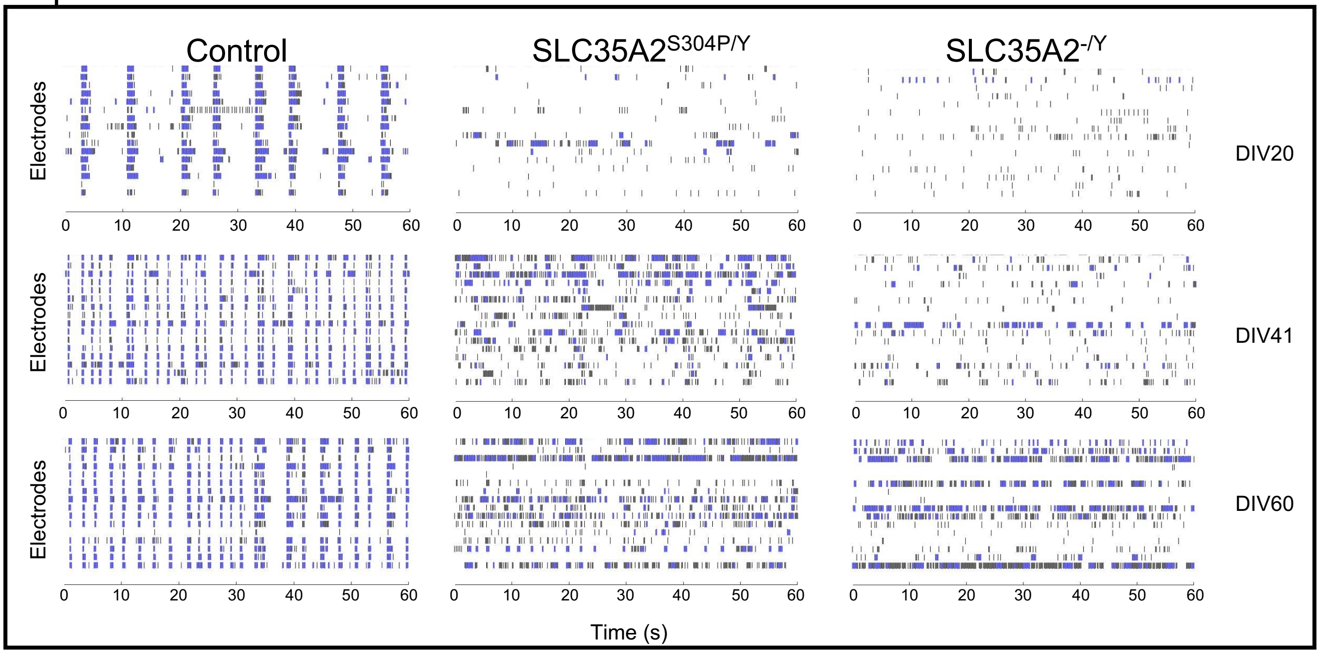


Experiment 4

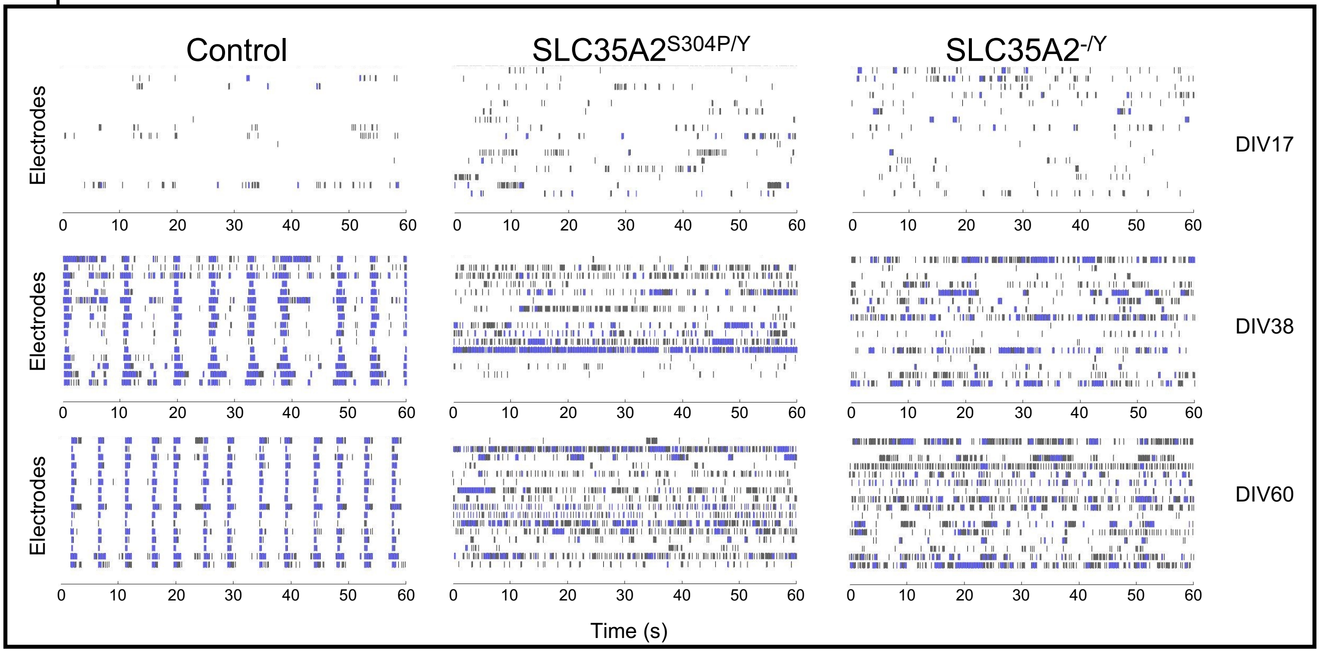


**Figure S9. Raster plots representative of each MEA plate throughout network development.** Raster plots capture electrode and network level activity. The activity of a single electrode is represented on the y-axis with each row representing an electrode. Spikes are detected as black ticks and bursts indicated in blue. The raster plots depicted here capture 60 seconds of the 900 second full length recording. Activity from each of the four MEA plates are represented as Experiment 1, Experiment 2, Experiment 3, and Experiment 4. The day *in vitro* (DIV) represents the number of days post-plating on MEA.

#### Supplementary Figure 10

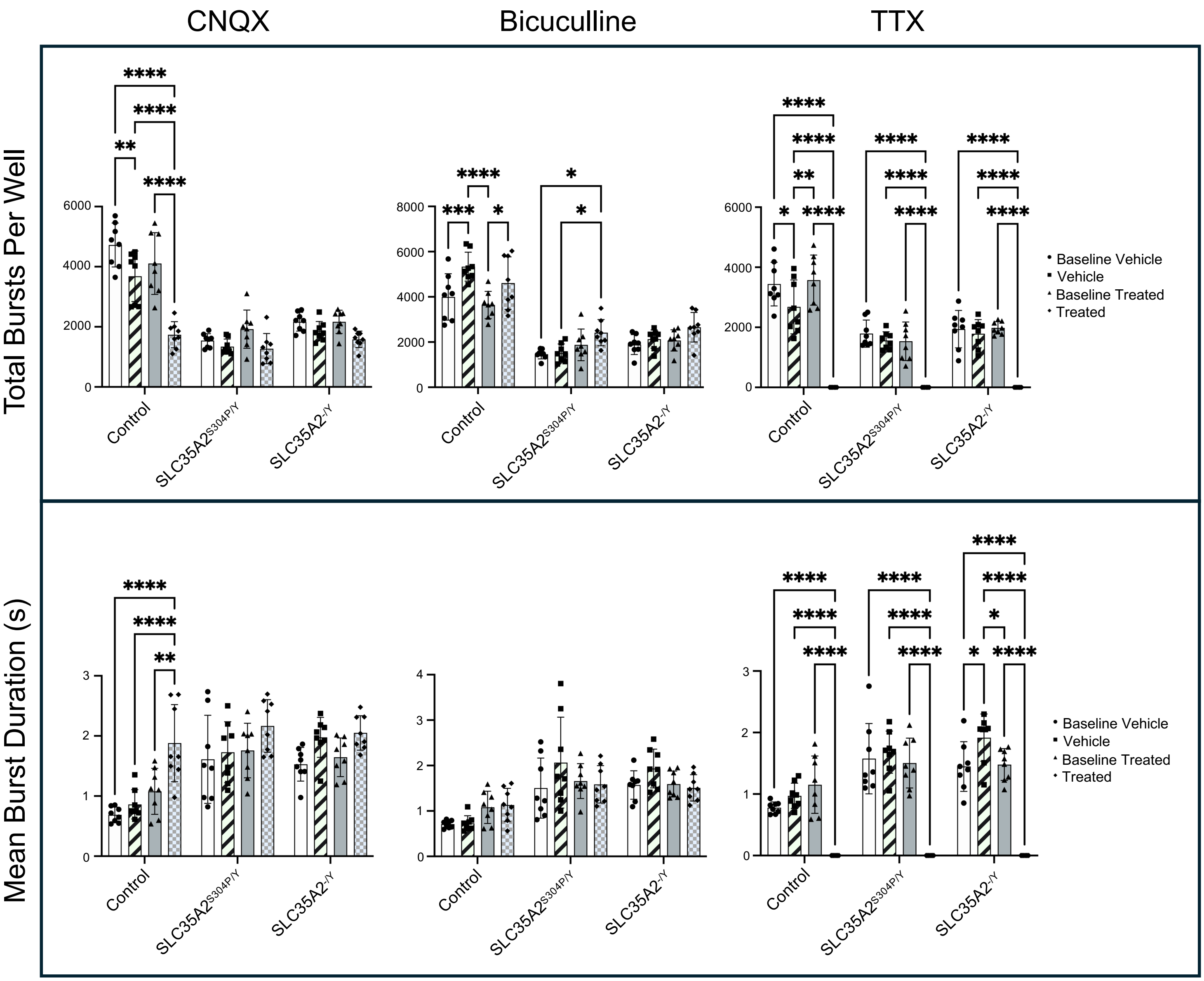


;

#### Supplementary Figure 10 (cont'd)

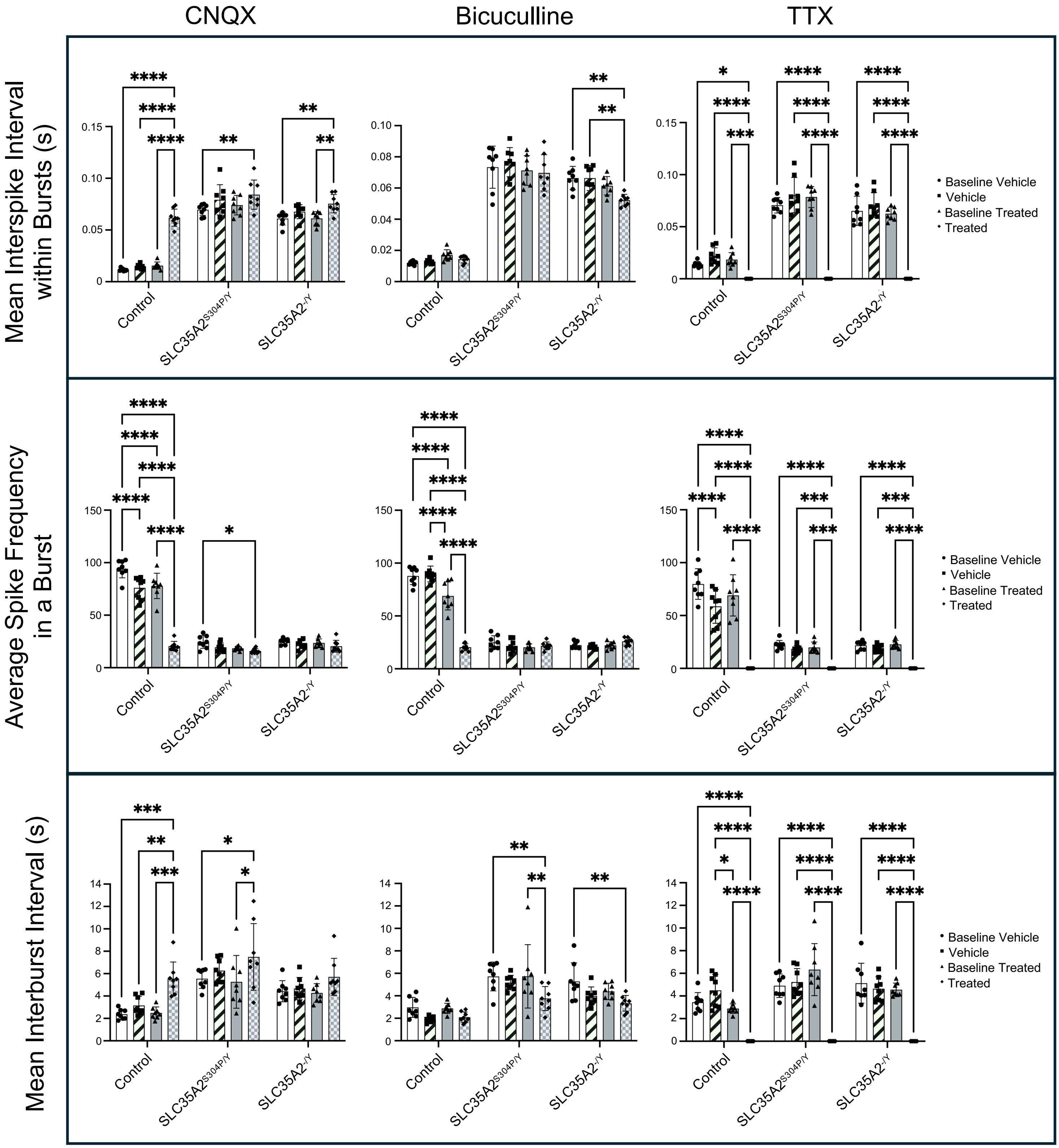

**Figure S10. Burst features of neural networks following pharmacologic perturbation.** Baseline activity of neural networks was recorded followed by treatment with vehicle or drug (CNQX, bicuculline, TTX). Treatment with each drug was carried out every 2-3 days after washout allowing networks to recover back to baseline. Data is representative of the mean and standard deviation of two pooled independent differentiations. Each dot represents data obtained from a single MEA well. Statistics: Two-way ANOVA test with Tukey’s to correct for multiple comparisons. * = $P \leq 0.05$; ** = $P \leq 0.01$; *** = $P \leq 0.001$ **** = $P \leq 0.0001$.

#### Supplementary Figure 11

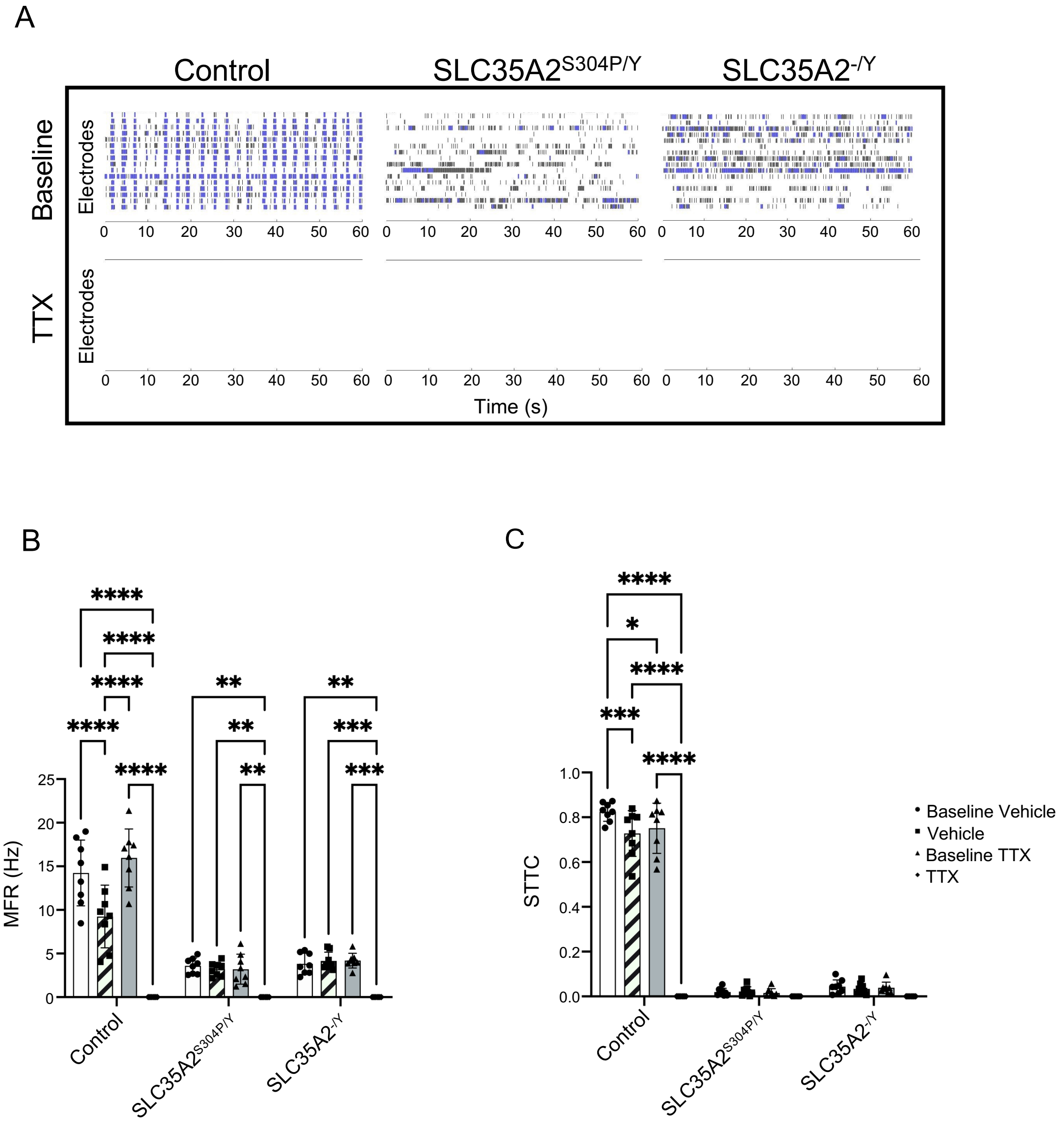


**Figure S11. Acute tetrodotoxin abolishes spontaneous network activity. (A)** Raster plots depicting baseline MEA recordings (top row) and after acute tetrodotoxin (TTX) treatment (bottom row). Raster plots represent a 60 sec snapshot of the 900 sec total recording time. **(B)** The effect of TTX on MFR by active electrode. **(C)** The effect of TTX on STTC. Data is representative of the mean and standard deviation of two pooled independent differentiations. Each dot represents data obtained from a single MEA well. Statistics: Two-way ANOVA with Tukey's multiple comparisons. Data is represented as mean and standard deviation. * = $P \leq 0.05$; ** = $P \leq 0.01$; *** = $P \leq 0.001$ **** = $P \leq 0.0001$.

#### Supplementary Figure 12

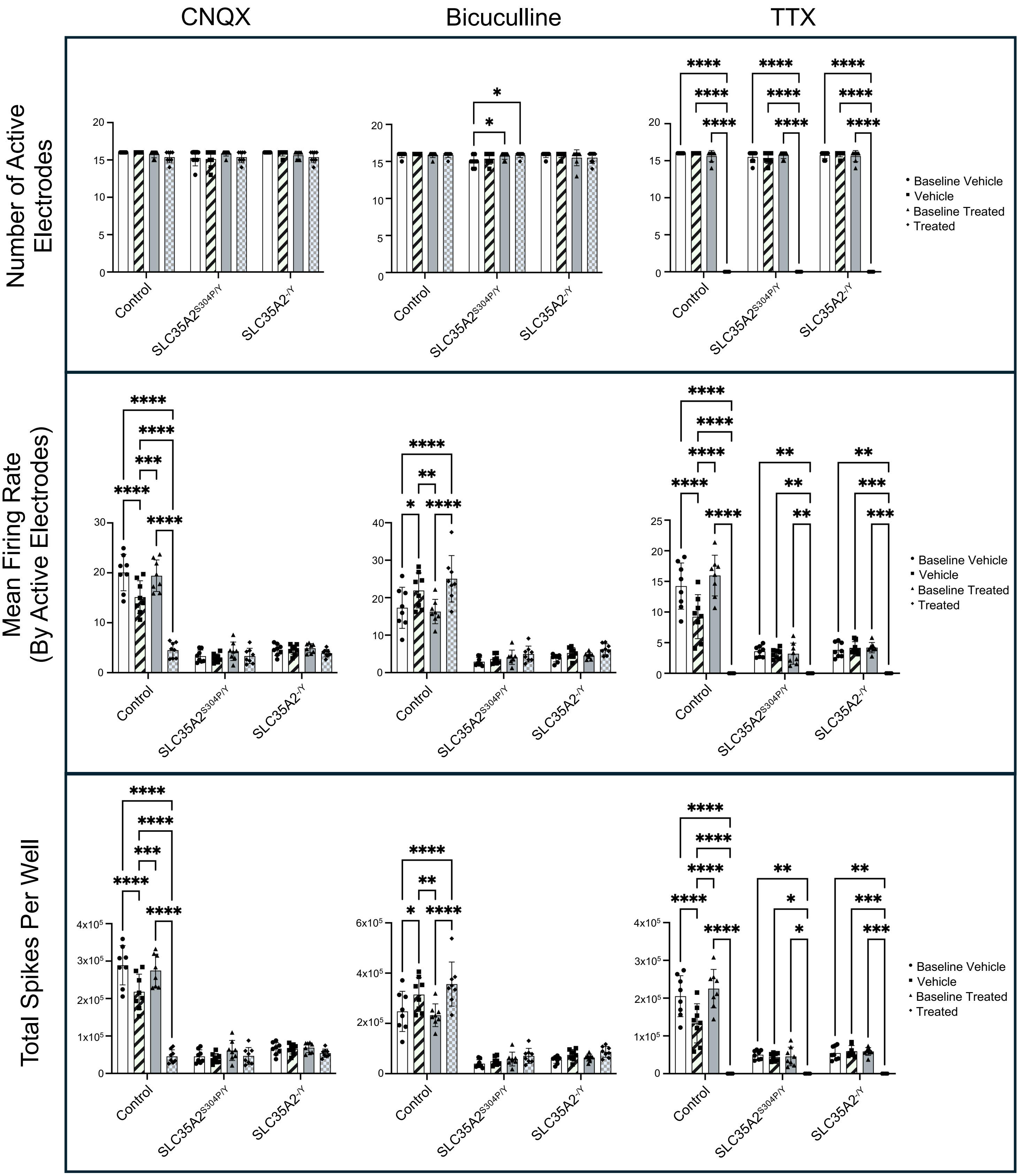


**Figure S12. Spike features of neural networks following pharmacologic perturbation.** Baseline activity of neural networks was recorded followed by treatment with vehicle or drug (CNQX, bicuculline, TTX). Treatment with each drug was carried out every 2-3 days after washout allowing networks to recover back to baseline. Data is representative of the mean and standard deviation of two pooled independent differentiations. Each dot represents data obtained from a single MEA well. Statistics: Two-way ANOVA test with Tukey's to correct for multiple comparisons. * = P≤0.05; ** = P≤0.01; *** = P≤0.001; **** = P≤0.0001.

#### Supplementary Figure 13

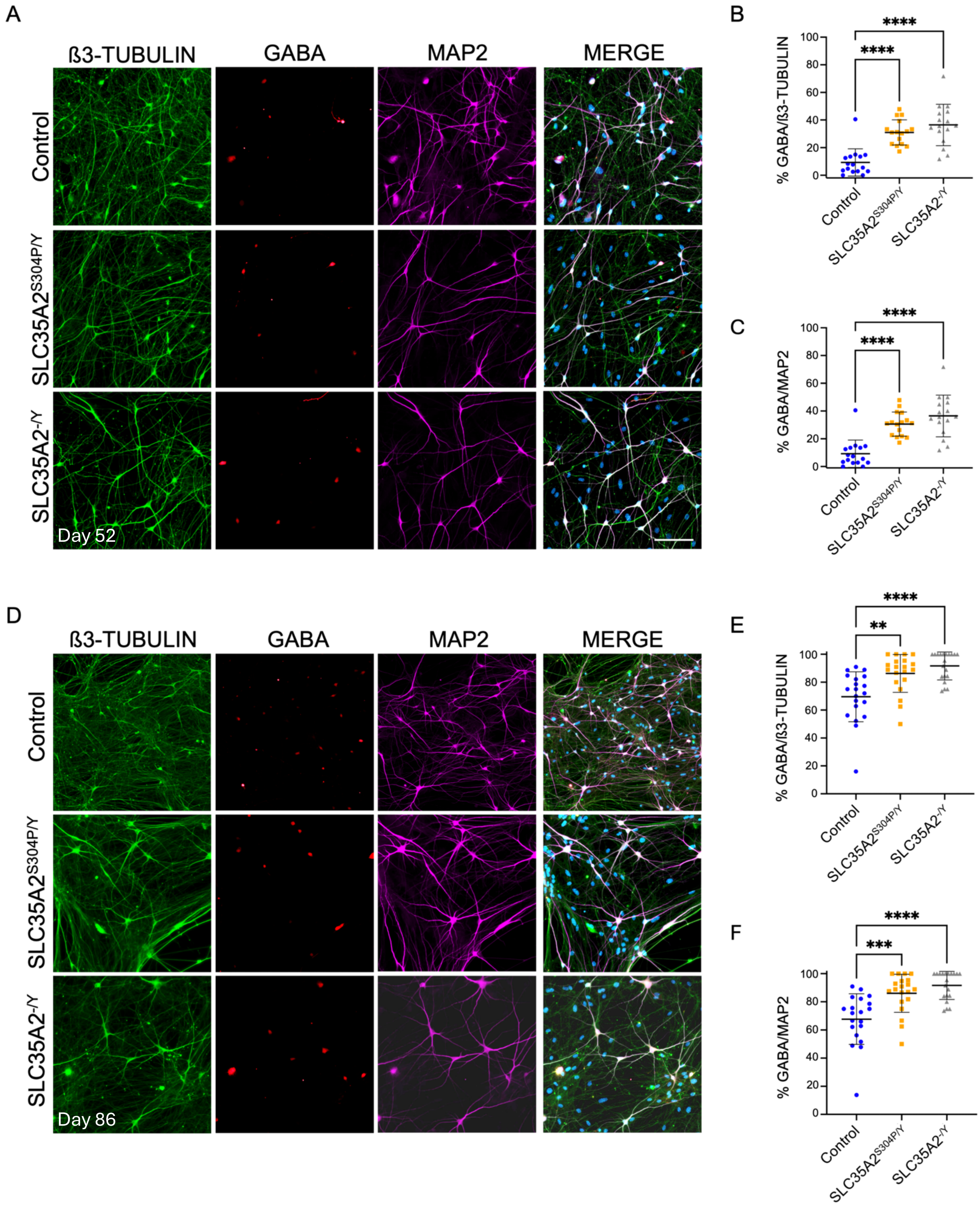


**Figure S13. SLC35A2$^{S304P/Y}$ and SLC35A2$^{-/Y}$ neurons influence neural trajectory towards the GABAergic fate. (A)** Immunofluorescent staining of GABAergic neurons in day 52 neuron cultures. Neurons were stained with ß3-TUBULIN (green), GABA (red), MAP2 (far red), and DAPI for nuclei. Quantification of **(B)** % GABA/ß3-TUBULIN and **(C)** % GABA/MAP2 in day 52 neuronal cultures. **(D)** Immunofluorescent staining of GABAergic neurons in day 86 neuron cultures. Neurons were stained with ß3-TUBULIN (green), GABA (red), MAP2 (far red), and DAPI for nuclei. Quantification of **(E)** % GABA/ß3-TUBULIN and **(F)** % GABA/MAP2 in day 86 neuronal cultures. Data is representative of two pooled independent differentiations. Statistics: One-way ANOVA test with Tukey's to correct for multiple comparisons. * = P≤0.05; ** = P≤0.01; *** = P≤0.001; **** = P≤0.0001.

#### Supplementary Figure 14

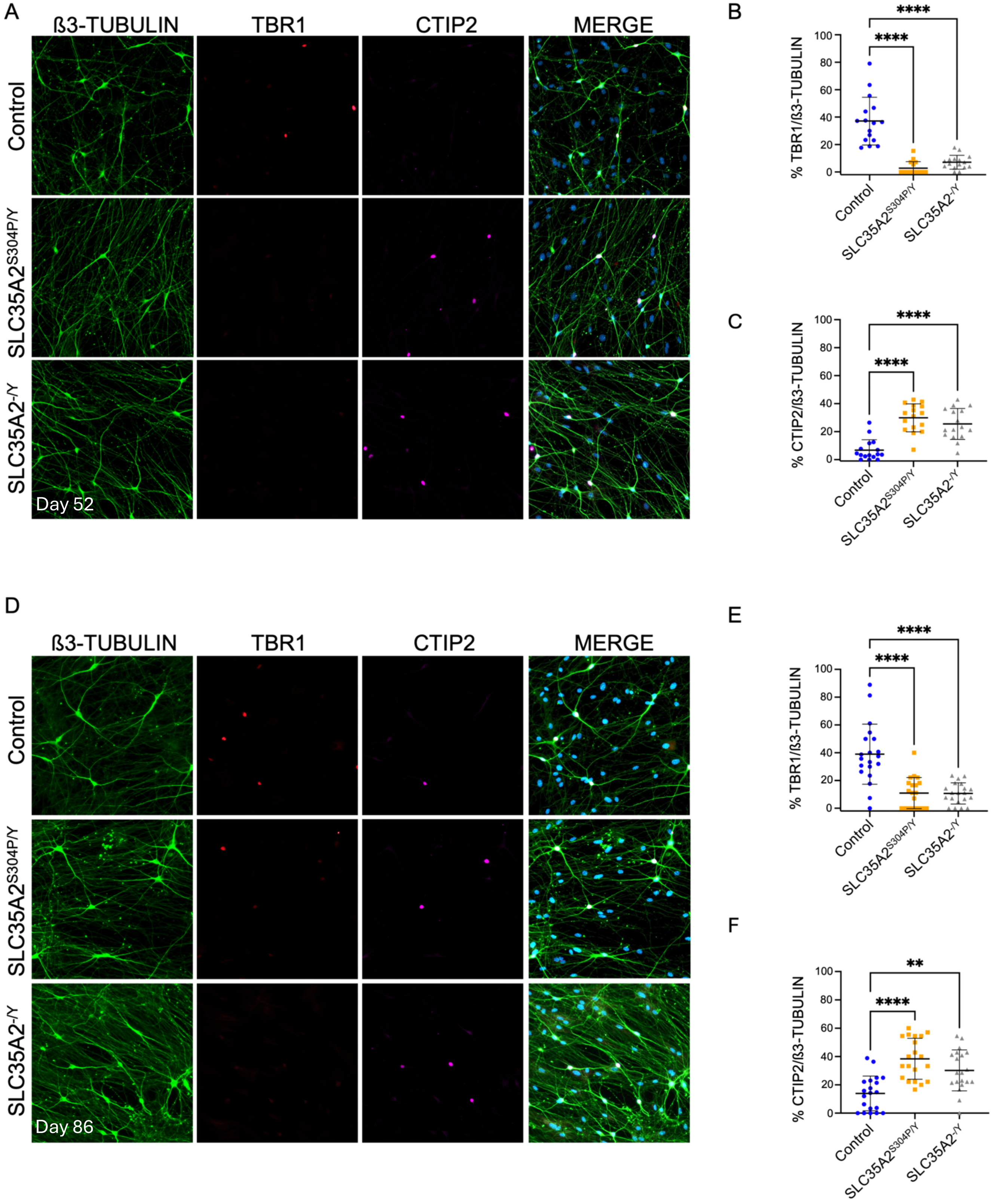


**Figure S14. Neural networks express deep layer cortical markers. (A)** Immunofluorescent staining of deep layer neurons in day 52 neuron cultures. Neurons were stained with ß3-TUBULIN (green), TBR1 (red), CTIP2 (far red), and DAPI for nuclei. Quantification of **(B)** % TBR1/ß3-TUBULIN and **(C)** % CTIP2/ß3-TUBULIN in day 52 neuronal cultures. **(D)** Immunofluorescent staining of deep layer neurons in day 86 neuron cultures. Neurons were stained with ß3-TUBULIN (green), TBR1 (red), CTIP2 (far red), and DAPI for nuclei. Quantification of **(E)** % TBR1/ß3-TUBULIN and **(F)** % CTIP2/ß3-TUBULIN in day 86 neuronal cultures. Data is representative of two pooled independent differentiations. Statistics: One-way ANOVA test with Tukey's to correct for multiple comparisons.

### Supplementary Figure 15

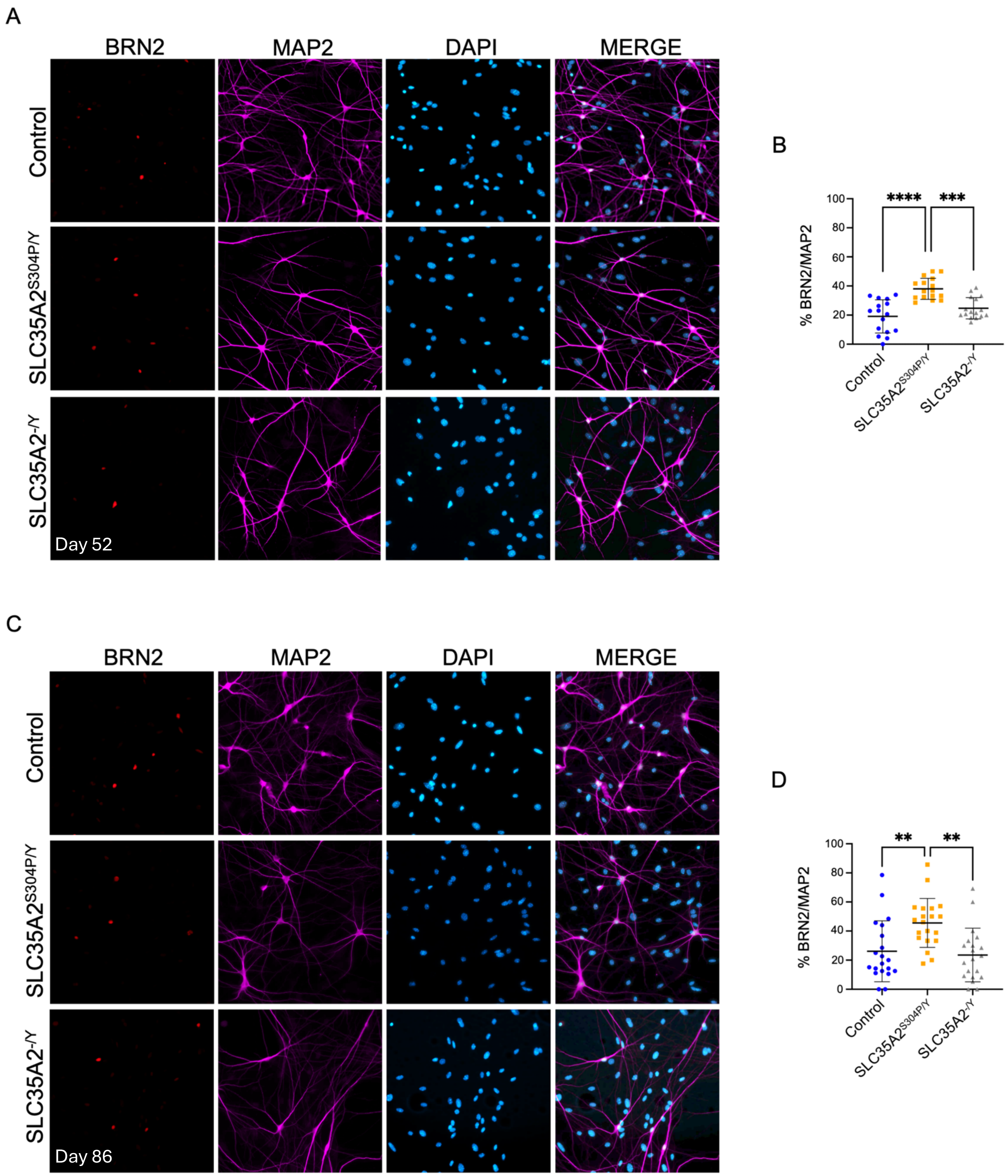


**Figure S15. Neural networks express upper layer cortical markers. (A)** Immunofluorescent staining of upper layer neurons in day 52 neuron cultures. Neurons were stained with BRN2 (red), MAP2 (far red), and DAPI for nuclei. **(B)** Quantification of % BRN2/MAP2. **(C)** Immunofluorescent staining of upper layer neurons in day 86 neuron cultures. Neurons were stained with BRN2 (red), MAP2 (far red), and DAPI for nuclei. **(D)** Quantification of % BRN2/MAP2 in day 86 neuronal cultures. Data is representative of two pooled independent differentiations. Statistics: One-way ANOVA test with Tukey's to correct for multiple comparisons. * = $P \leq 0.05$; ** = $P \leq 0.01$; *** = $P \leq 0.001$; **** = $P \leq 0.0001$.

### Supplementary Figure 16

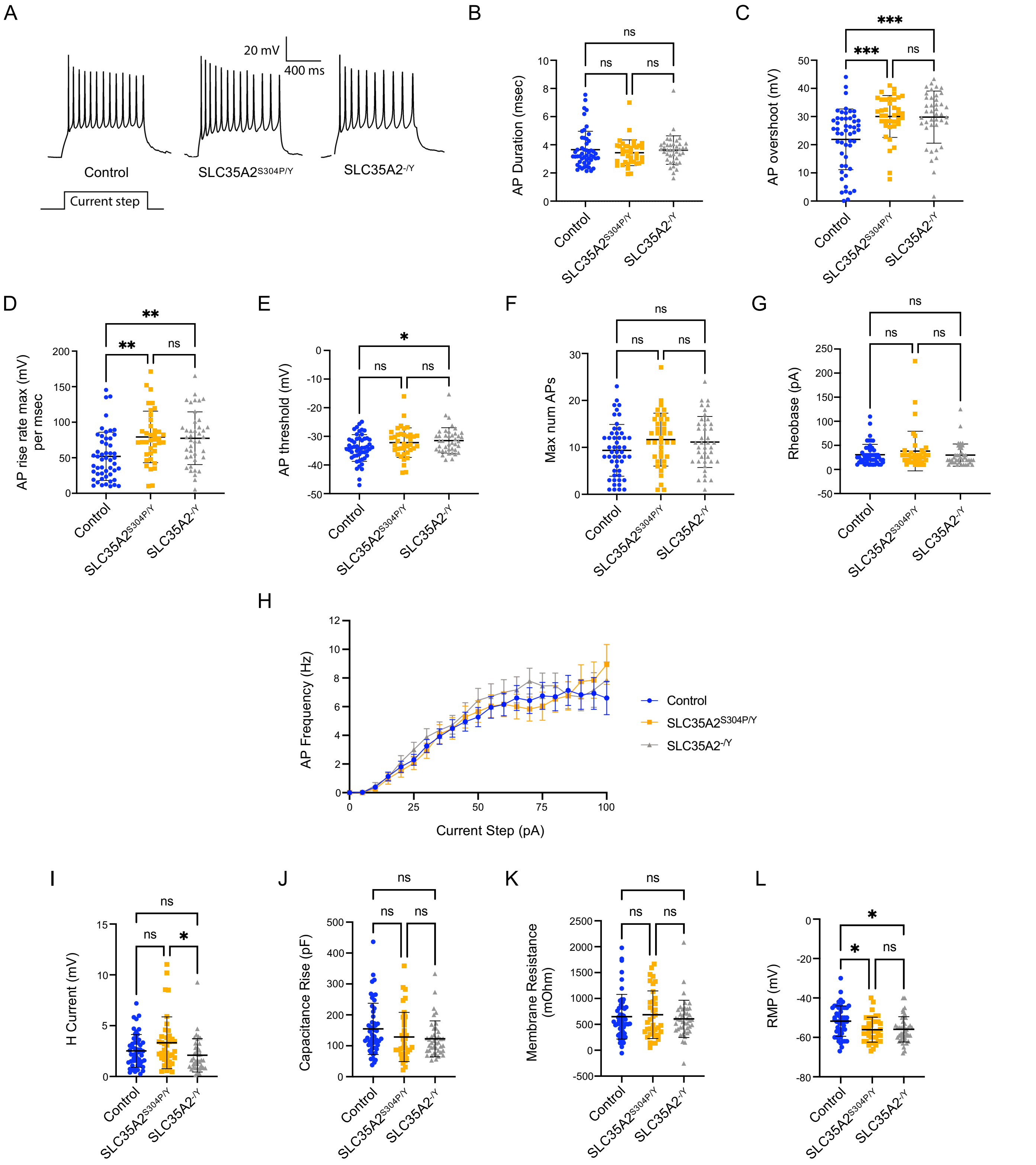


**Figure S16. Whole-cell current clamp recordings reveal no differences consistent with intrinsic excitability changes.** Action potential (AP) properties were assessed in figures (A) through (H). **(A)** Membrane voltage traces showing AP firing in response to a 1 s depolarizing current step of the amplitude which evokes the maximum number of APs in that cell. **(B)** AP duration (msec). **(C)** AP overshoot (mV). **(D)** Maximum AP rise rate (ms). **(E)** AP threshold (mV). **(F)** Maximum number of APs in a 1 s current step. **(G)** Rheobase (pA). **(H)** AP frequency in response to current steps increasing in 5 pA increments. Passive membrane properties were assessed in figures (I) through (L). **(I)** H current (mV). **(J)** Capacitance rise (pF). **(K)** Membrane resistance (MOhm). **(L)** Resting membrane potential (RMP, mV). The data is representative of five pooled independent differentiations with each dot representing recordings from a single neuron. Statistics: One-way ANOVA test with Tukey's to correct for multiple comparisons. * = P≤0.05; ** = P≤0.01; *** = P≤0.001 **** = P≤0.0001.

### Supplementary Table 1.

**Table S1.** List of antibodies used for immunofluorescent staining.

| Antibody | Species | Dilution | Manufacturer | Catalog # |
|---|---|---|---|---|
| SLC35A2 | Rabbit | 1:50 | Novus Biologicals | NBP1-80642 |
| GM-130 | Mouse | 1:200 | Novus Biologicals | H00002801-B01P |
| OCT3/4 | Mouse | 1:200 | Santa Cruz | sc-5279 |
| NANOG | Rabbit | 1:100 | Abcam | ab21624 |
| TRA-1-81 | Mouse | 1:100 | Stemgent | 09-0011 |
| SOX2 | Rabbit | 1:100 | Cell Signaling Technology | 3579 |
| NESTIN | Mouse | 1:1000 | Stem Cell Technologies | 60091.1 |
| PAX6 | Mouse | 1:200 | DSHB | PAX6-b |
| PAX6 | Rabbit | 1:100 | BioLegend | PRB-278P |
| ZO-1 | Rat | 1:100 | Santa Cruz | sc-33725 |
| ZO-1 | Mouse | 1:100 | Invitrogen | 339100 |
| ß3-TUBULIN | Rabbit | 1:1000 | BioLegend | 802001 |
| MAP2 | Mouse | 1:500 | Sigma Aldrich | M4403 |
| MAP2 | Chicken | 1:10,000 | BioLegend | PCK-554P |
| GABA | Rabbit | 1:200 | Thermo Scientific | PA5-32241 |
| TBR1 | Rabbit | 1:1000 | Abcam | ab31940 |
| CTIP2 | Rat | 1:600 | Abcam | ab18465 |
| BRN2 | Rabbit | 1:250 | Abcam | ab137469 |
| Goat Anti-Mouse Alexa Fluor 488 | | 1:1000 | Invitrogen | A32723 |
| Goat Anti-Rabbit Alexa Fluor 488 | | 1:1000 | Invitrogen | A11034 |
| Goat Anti-Mouse Alexa Fluor 568 | | 1:1000 | Invitrogen | A21043 |
| Goat Anti-Rabbit Alexa Fluor 568 | | 1:1000 | Invitrogen | A11036 |
| Goat Anti-Rat Alexa Fluor 647 | | 1:1000 | Invitrogen | A21247 |
| Goat Anti-Chicken Alexa Fluor 647 | | 1:1000 | Invitrogen | A21449 |
| Goat Anti-Guinea Pig Alexa Fluor 647 | | 1:1000 | Invitrogen | A21450 |

### Supplementary Table 2.

**Table S2.** List of antibodies used for Western blot.

| Antibody | Species | Dilution | Manufacturer | Catalog # |
|---|---|---|---|---|
| SLC35A2 | Rabbit | 1:500 | Novus Biologicals | NBP1-80642 |
| TRA-1-81 | Mouse | 1:1000 | Stemgent | 09-0011 |
| Podocalyxin | Mouse | 1:500 | Santa Cruz Biotechnology | sc-23904 |
| GAPDH | Rabbit | 1:1000 | Cell Signaling Technology | 2118S |
| Goat Anti-Mouse IgG (H+L), HRP | | 1:10,000 | Invitrogen | G21040 |
| Goat Anti-Rabbit IgG (H+L), HRP | | 1:10,000 | Invitrogen | G21234 |

#### Supplementary Table 3.

**Table S3.** Separation gradient method.

| Time [min] | Flow [mL/min] | A% | B% |
|---|---|---|---|
| 0 | 0.5 | 20 | 80 |
| 2 | 0.5 | 25 | 75 |
| 48 | 0.5 | 38 | 62 |
| 49 | 0.5 | 60 | 40 |
| 51.5 | 0.5 | 20 | 80 |
| 52 | 0.5 | 20 | 80 |
| 60 | 0.5 | 20 | 80 |

### Supplementary Table 4.

**Table S4.** List of Taqman assays used for qRT-PCR.

| Taqman | Assay ID |
|---|---|
| SLC35A2 | Hs00194644_m1 |
| NANOG | Hs02387400_g1 |
| POU5F1 | Hs00999632_g1 |
| SOX2 | Hs01053049_s1 |
| SLC32A1 | Hs00369773_m1 |
| SLC6A1 | Hs01104469_m1 |
| SLC17A7 | Hs01574214_m1 |
| GAD1 | Hs01065887_m1 |
| GLS | Hs01014020_m1 |
| DLX2 | Hs00269993_m1 |
| DLX5 | Hs00193291_m1 |
| DLX6 | Hs00231999_m1 |
| LHX6 | Hs01030943_m1 |
| NKX2.1 | Hs03987940_m1 |
| NR2F2 | Hs01041380_g1 |
| SYN1 | Hs00199577_m1 |
| DLG4 | Hs01555373_m1 |
| GPHN | Hs00982840_m1 |
| FOXG1 | Hs01850784_s1 |
| EMX1 | Hs05360123_s1 |
| GSX2 | Hs00370195_m1 |
| DARPP32 | Hs00259967_m1 |
| GAPDH | Hs99999905_m1 |

### Supplementary Table 5

**Table S5.** Number of wells plated per genotype for each MEA experiment. Four biologically independent differentiations were plated onto individual MEA plates (denoted Experiments 1 through 4). Technical replicates, which represent differentiations initiated at the same time point from independent wells are designated as A and B. Astrocyte only wells were also included on MEA plates to ensure no contaminating mouse neurons are contributing to measured activity.

| iPSC Line | Experiment 1-A | Experiment 1-B | Experiment 2-A | Experiment 2-B | Experiment 3 | Experiment 4-A | Experiment 4-B | Total Wells |
|---|---|---|---|---|---|---|---|---|
| WT | 7 | 7 | 10 | 6 | 7 | 8 | 8 | 53 |
| S304P | 7 | 7 | 6 | 7 | 7 | 8 | 8 | 50 |
| Indel | 7 | 7 | 7 | 7 | 7 | 8 | 8 | 51 |

### Supplementary Table 6

**Table S6.** A statistical summary of MEA spike features. SLC35A2$^{S304P/Y}$ and SLC35A2$^{-/Y}$ neural networks share similar activity patterns. The "direction" of change for each feature per cell line is shown relative to the control. The displayed permutation p-value (perm_pval) represents the p-value derived using a Mann Whitney test followed by 1000 permutations. The p-value for each experimental replicate is shown per feature relative to the control.

| Feature | Genotype | | Experiment 1A | Experiment 1B | Experiment 2A | Experiment 2B | Experiment 3 | Experiment 4A | Experiment 4B |
|---|---|---|---|---|---|---|---|---|---|
| Number of Active Electrodes | SLC35A2$^{S304P/Y}$ | perm_pval | 0.196 | 0.363 | <0.001 | 0.123 | 0.195 | 0.002 | 0.112 |
| | | Direction | Same | Same | Slight Decrease | Same | Same | Slight Decrease | Same |
| | SLC35A2$^{-/Y}$ | perm_pval | 0.575 | 0.453 | 0.005 | 0.17 | 0.032 | 0.001 | 0.552 |
| | | Direction | Same | Same | Slight Decrease | Same | Slight Decrease | Slight Decrease | Same |
| Mean Firing Rate Across Active Electrodes | SLC35A2$^{S304P/Y}$ | perm_pval | <0.001 | <0.001 | <0.001 | <0.001 | <0.001 | <0.001 | <0.001 |
| | | Direction | Decrease | Decrease | Decrease | Decrease | Decrease | Decrease | Decrease |
| | SLC35A2$^{-/Y}$ | perm_pval | <0.001 | <0.001 | 0.01 | <0.001 | <0.001 | <0.001 | <0.001 |
| | | Direction | Decrease | Decrease | Slight Decrease | Decrease | Decrease | Decrease | Decrease |
| STTC | SLC35A2$^{S304P/Y}$ | perm_pval | <0.001 | <0.001 | <0.001 | <0.001 | <0.001 | <0.001 | <0.001 |
| | | Direction | Decrease | Decrease | Decrease | Decrease | Decrease | Decrease | Decrease |
| | SLC35A2$^{-/Y}$ | perm_pval | <0.001 | <0.001 | <0.001 | <0.001 | <0.001 | <0.001 | <0.001 |
| | | Direction | Decrease | Decrease | Decrease | Decrease | Decrease | Decrease | Decrease |
| Mutual Information | SLC35A2$^{S304P/Y}$ | perm_pval | <0.001 | <0.001 | <0.001 | <0.001 | <0.001 | <0.001 | <0.001 |
| | | Direction | Decrease | Decrease | Decrease | Decrease | Decrease | Decrease | Decrease |
| | SLC35A2$^{-/Y}$ | perm_pval | <0.001 | <0.001 | <0.001 | <0.001 | <0.001 | <0.001 | <0.001 |
| | | Direction | Decrease | Decrease | Decrease | Decrease | Decrease | Decrease | Decrease |

#### Supplementary Table 7

**Table S7.** A statistical summary of MEA network spike features. SLC35A2$^{S304P/Y}$ and SLC35A2$^{-/Y}$ neural networks share similar activity patterns. The "direction" of change for each feature per cell line is shown relative to the control. The displayed permutation p-value (perm_pval) represents the p-value derived using a Mann Whitney test followed by 1000 permutations. The p-value for each experimental replicate is shown per feature relative to the control.

| Feature | Genotype | | Experiment 1A | Experiment 1B | Experiment 2A | Experiment 2B | Experiment 3 | Experiment 4A | Experiment 4B |
|---|---|---|---|---|---|---|---|---|---|
| Network Spikes | SLC35A2$^{S304P/Y}$ | perm_pval | <0.001 | <0.001 | <0.001 | <0.001 | <0.001 | <0.001 | <0.001 |
| | | Direction | Decrease | Decrease | Decrease | Decrease | Decrease | Decrease | Decrease |
| | SLC35A2$^{-/Y}$ | perm_pval | <0.001 | <0.001 | <0.001 | <0.001 | <0.001 | <0.001 | <0.001 |
| | | Direction | Decrease | Decrease | Decrease | Decrease | Decrease | Decrease | Decrease |
| % Spikes in Network Spikes | SLC35A2$^{S304P/Y}$ | perm_pval | <0.001 | <0.001 | <0.001 | <0.001 | <0.001 | <0.001 | <0.001 |
| | | Direction | Decrease | Decrease | Decrease | Decrease | Decrease | Decrease | Decrease |
| | SLC35A2$^{-/Y}$ | perm_pval | <0.001 | <0.001 | <0.001 | <0.001 | <0.001 | <0.001 | <0.001 |
| | | Direction | Decrease | Decrease | Decrease | Decrease | Decrease | Decrease | Decrease |
| Mean Spikes in Network Spikes | SLC35A2$^{S304P/Y}$ | perm_pval | <0.001 | <0.001 | <0.001 | <0.001 | <0.001 | <0.001 | <0.001 |
| | | Direction | Decrease | Decrease | Decrease | Decrease | Decrease | Decrease | Decrease |
| | SLC35A2$^{-/Y}$ | perm_pval | <0.001 | <0.001 | <0.001 | <0.001 | <0.001 | <0.001 | <0.001 |
| | | Direction | Decrease | Decrease | Decrease | Decrease | Decrease | Decrease | Decrease |

### Supplementary Table 8

**Table S8.** A statistical summary of MEA burst features. SLC35A2$^{S304P/Y}$ and SLC35A2$^{-/Y}$ neural networks share similar activity patterns. The "direction" of change for each feature per cell line is shown relative to the control. The displayed permutation p-value (perm_pval) represents the p-value derived using a Mann Whitney test followed by 1000 permutations. The p-value for each experimental replicate is shown per feature relative to the control.

| Feature | Genotype | | Experiment 1A | Experiment 1B | Experiment 2A | Experiment 2B | Experiment 3 | Experiment 4A | Experiment 4B |
|---|---|---|---|---|---|---|---|---|---|
| Mean Burst Duration | SLC35A2$^{S304P/Y}$ | perm_pval | <0.001 | <0.001 | <0.001 | 0.001 | <0.001 | 0.031 | 0.024 |
| | | Direction | Increase | Increase | Increase | Increase | Increase | Increase | Slight Increase |
| | SLC35A2$^{-/Y}$ | perm_pval | <0.001 | <0.001 | <0.001 | <0.001 | 0.009 | <0.001 | 0.106 |
| | | Direction | Increase | Increase | Increase | Increase | Increase | Increase | Slight Increase |
| Number of Bursts Per Well | SLC35A2$^{S304P/Y}$ | perm_pval | <0.001 | <0.001 | <0.001 | <0.001 | 0.001 | <0.001 | <0.001 |
| | | Direction | Decrease | Decrease | Decrease | Decrease | Decrease | Decrease | Decrease |
| | SLC35A2$^{-/Y}$ | perm_pval | <0.001 | <0.001 | 0.002 | <0.001 | 0.003 | <0.001 | <0.001 |
| | | Direction | Decrease | Decrease | Decrease | Decrease | Decrease | Decrease | Decrease |
| Bursts Per Minute | SLC35A2$^{S304P/Y}$ | perm_pval | <0.001 | <0.001 | <0.001 | <0.001 | <0.001 | <0.001 | <0.001 |
| | | Direction | Decrease | Decrease | Decrease | Decrease | Decrease | Decrease | Decrease |
| | SLC35A2$^{-/Y}$ | perm_pval | <0.001 | <0.001 | 0.002 | <0.001 | 0.001 | <0.001 | <0.001 |
| | | Direction | Decrease | Decrease | Decrease | Decrease | Decrease | Decrease | Decrease |
| Mean Spike Frequency in Bursts | SLC35A2$^{S304P/Y}$ | perm_pval | <0.001 | <0.001 | <0.001 | <0.001 | <0.001 | <0.001 | <0.001 |
| | | Direction | Decrease | Decrease | Decrease | Decrease | Decrease | Decrease | Decrease |
| | SLC35A2$^{-/Y}$ | perm_pval | <0.001 | <0.001 | <0.001 | <0.001 | <0.001 | <0.001 | <0.001 |
| | | Direction | Decrease | Decrease | Decrease | Decrease | Decrease | Decrease | Decrease |
| Mean Spikes in Bursts | SLC35A2$^{S304P/Y}$ | perm_pval | <0.001 | <0.001 | 0.004 | 0.003 | 0.012 | <0.001 | <0.001 |
| | | Direction | Decrease | Decrease | Slight Decrease | Slight Decrease | Slight Decrease | Decrease | Decrease |
| | SLC35A2$^{-/Y}$ | perm_pval | 0.006 | <0.001 | 0.446 | 0.002 | 0.018 | <0.001 | <0.001 |
| | | Direction | Slight Decrease | Decrease | Same | Slight Decrease | Slight Decrease | Decrease | Decrease |
| Mean Inter-Burst Interval | SLC35A2$^{S304P/Y}$ | perm_pval | <0.001 | <0.001 | <0.001 | <0.001 | <0.001 | <0.001 | 0.001 |
| | | Direction | Slight Increase | Slight Increase | Slight Increase | Slight Increase | Slight Increase | Slight Increase | Slight Increase |
| | SLC35A2$^{-/Y}$ | perm_pval | 0.004 | 0.033 | 0.001 | <0.001 | <0.001 | <0.001 | <0.001 |
| | | Direction | Slight Increase | Slight Increase | Slight Increase | Slight Increase | Slight Increase | Slight Increase | Slight Increase |
| Percent Spikes in Burst | SLC35A2$^{S304P/Y}$ | perm_pval | <0.001 | <0.001 | <0.001 | <0.001 | <0.001 | <0.001 | <0.001 |
| | | Direction | Decrease | Decrease | Decrease | Decrease | Decrease | Decrease | Decrease |
| | SLC35A2$^{-/Y}$ | perm_pval | <0.001 | <0.001 | <0.001 | <0.001 | <0.001 | <0.001 | <0.001 |
| | | Direction | Decrease | Decrease | Decrease | Decrease | Decrease | Decrease | Decrease |
| Mean Inter-Spike Interval within Bursts | SLC35A2$^{S304P/Y}$ | perm_pval | <0.001 | <0.001 | <0.001 | <0.001 | <0.001 | <0.001 | <0.001 |
| | | Direction | Increase | Increase | Increase | Increase | Increase | Increase | Increase |
| | SLC35A2$^{-/Y}$ | perm_pval | <0.001 | <0.001 | <0.001 | <0.001 | <0.001 | <0.001 | <0.001 |
| | | Direction | Increase | Increase | Increase | Increase | Increase | Increase | Increase |

### Supplementary Blots

A

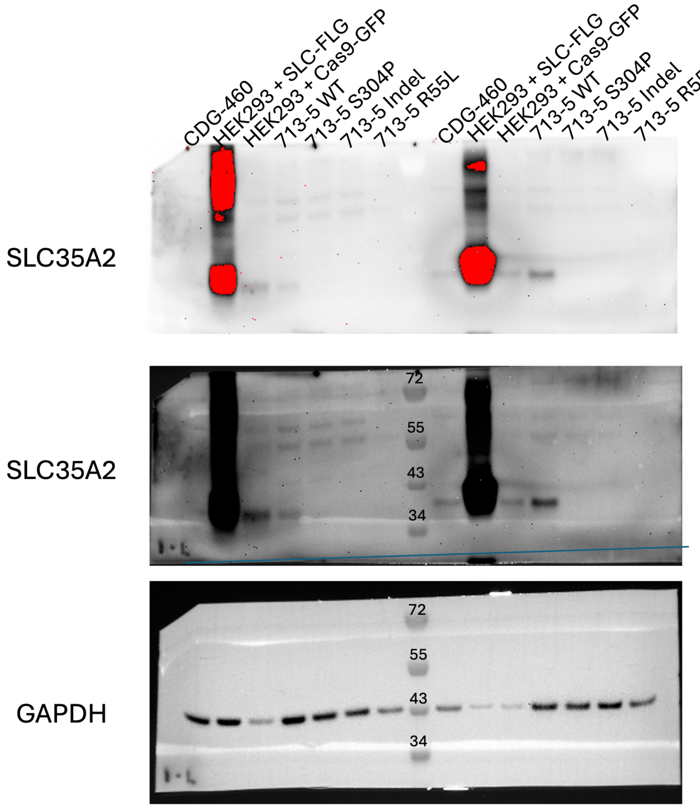


B

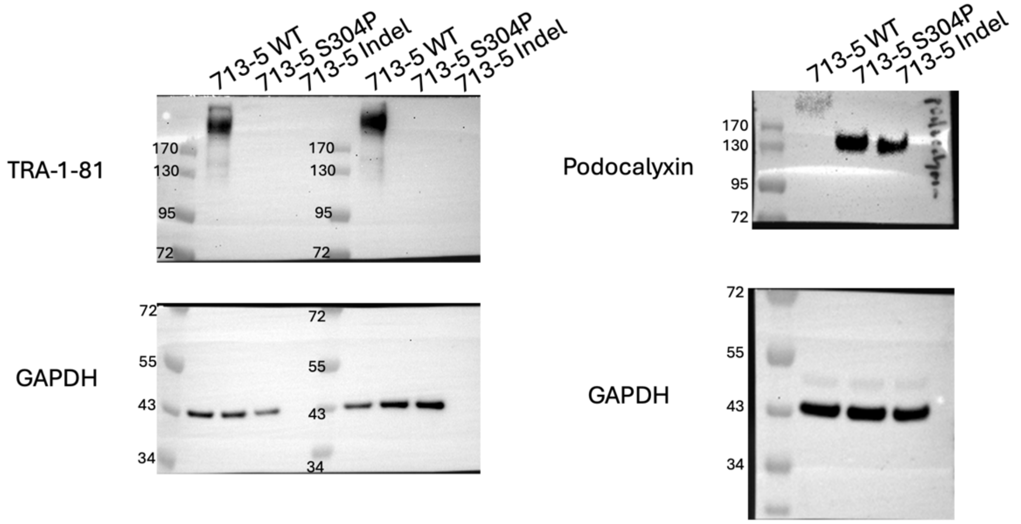


**Supplementary Blots.** Representative full-length western blots from Fig. 1C and Supplementary Figure 1E. **(A)** Membranes were probed with an SLC35A2 antibody and GAPDH as a loading control. The antibody used in this study has been previously published (Sosicka et al, 2019). HEK293 transfected with an SLC35A2 plasmid was used as a positive control (HEK293 + SLC-FLG). **(B)** Membranes were probed with a TRA1-81 and podocalyxin antibody with GAPDH as a loading control.
